# Post-vaccination Omicron infections induce broader immunity across antigenic space than prototype mRNA COVID-19 booster vaccination or primary infection

**DOI:** 10.1101/2022.07.05.498883

**Authors:** Wei Wang, Sabrina Lusvarghi, Rahul Subramanian, Nusrat J. Epsi, Richard Wang, Emilie Goguet, Anthony C Fries, Fernando Echegaray, Russell Vassell, Si’Ana Coggins, Stephanie A. Richard, David A. Lindholm, Katrin Mende, Evan Ewers, Derek Larson, Rhonda E. Colombo, Christopher Colombo, Janet O. Joseph, Julia Rozman, Alfred Smith, Tahaniyat Lalani, Catherine Berjohn, Ryan Maves, Milissa Jones, Rupal Mody, Nikhil Huprikar, Jeffrey Livezey, David Saunders, Monique Hollis-Perry, Gregory Wang, Anuradha Ganesan, Mark P. Simons, Christopher C. Broder, David Tribble, Eric D. Laing, Brian Agan, Timothy H. Burgess, Edward Mitre, Simon D. Pollett, Leah C. Katzelnick, Carol D. Weiss

## Abstract

The rapid emergence of new SARS-CoV-2 variants challenges vaccination strategies. Here, we measured antigenic diversity among variants and interpreted neutralizing antibody responses following single and multiple exposures in longitudinal infection and vaccine cohorts. Antigenic cartography using primary infection antisera showed that BA.2, BA.4/BA.5, and BA.2.12.1 are distinct from BA.1 and closer to the Beta cluster. Three doses of an mRNA COVID-19 vaccine increased breadth to BA.1 more than to BA.4/BA.5 or BA.2.12.1. Omicron BA.1 post-vaccination infection elicited antibody landscapes characterized by broader immunity across antigenic space than three doses alone, although with less breadth than expected to BA.2.12.1 and BA.4/BA.5. Those with Omicron BA.1 infection after two or three vaccinations had similar neutralizing titer magnitude and antigenic breadth. Accounting for antigenic differences among variants of concern when interpreting neutralizing antibody titers aids understanding of complex patterns in humoral immunity and informs selection of future COVID-19 vaccine strains.

## INTRODUCTION

There is an urgent need to develop vaccination strategies to provide the broadest immunity against emerging and yet-to-emerge SARS-CoV-2 variants. COVID-19 has resulted in over 6.3 million deaths and 540 million infections worldwide (World Health Organization, 2020). SARS-CoV-2 continues to circulate globally, even as population immunity continues to increase due to infections, reinfections, primary series vaccination and/or vaccine boosting (Bergeri et al., 2022). While authorized and licensed COVID-19 vaccines provide substantial protection against severe COVID-19, new emerging SARS-CoV-2 variants continue to threaten their effectiveness, even after vaccine boosting. An increased reinfection risk associated with the Omicron variant compared to earlier SARS-CoV-2 variants has been observed (Pulliam et al., 2022). Approved or authorized COVID-19 vaccines encode the spike protein of first SARS-CoV-2 strain to emerge, Wuhan-Hu-1, defined as the ancestral strain. An antigenically divergent strain, Omicron (BA.1), was first identified in November 2021 and has led to millions of infections, including post-vaccine infections (PVI), and prompting further recommendations for boosting. Additional variants closely related to Omicron, including BA.2 and its descendants were detected soon afterwards. Strikingly, these have rapidly outcompeted BA.1 strains. For example, BA.2.12.1, and BA.4 and BA.5 are now collectively now the most common variants in the United States (Center for Disease Control, 2022; UK Health Security Agency, 2022; Wang et al., 2022).

Vaccine formulations based on the ancestral Wuhan-Hu-1 strain antigen continue to be used for both primary series and booster vaccination schedules (World Health Organization, 2022b). A critical public health question is whether vaccinations derived from more recent strains substantially increase immune magnitude and breadth above boosting with the same ancestral strain, including in populations which may be unvaccinated, vaccinated, boosted, infected, reinfected, or various combinations thereof. It is known that three doses of COVID-19 mRNA vaccines containing the ancestral strain broaden immunity against a range of variants (Lusvarghi et al., 2022). However, fourth doses with the ancestral strain only transiently boost neutralizing antibody titers back to the peak observed after three (Bar-On et al., 2022; Magen et al., 2022; Regev-Yochay et al., 2022). In contrast, sequential exposure to the ancestral vaccine followed by Omicron PVI may induce broader neutralizing antibody responses than vaccination with three doses alone (Quandt et al., 2022) although other studies suggest protection against severe disease is similar (Gagne et al., 2022).

Optimal timing and composition of SARS-CoV-2 vaccines for both boosters and primary series therefore remain unclear. The World Health Organization (WHO) recently noted that an Omicron vaccine may provide broader protection against emerging variants in individuals who have already received two doses of ancestral vaccines. WHO recommended that individuals who have not received a primary vaccine dose should still receive at least two doses of the ancestral-based vaccine rather than a single Omicron-based vaccine alone (World Health Organization, 2022a). Recently released preliminary results involving bivalent vaccines containing both the ancestral strain and Omicron BA.1 suggest that they induce similar or broader immunity against BA.1 than a third dose with the ancestral strain alone (Chalkias et al., 2022; Pfizer, 2022).

Antigenic diversity between Omicron variants has further complicated vaccine composition decision making. For example, a BA.1 booster may not provide sufficiently broad protection if more recently emerged variants like BA.2.12.1 and BA.4/BA.5 escape immunity more than BA.1. BA.2.12.1 and BA.4/BA.5 contain additional spike mutations that make them more resistant than BA.1 or BA.2 to neutralization by sera from individuals with three vaccine doses (Hachmann et al., 2022; Qu et al., 2022; Wang *et al*., 2022). Furthermore, individuals who have been vaccinated with BNT162b2 or vaccinated and infected with BA.1 or BA.2 have lower neutralizing antibody titers against BA.2.12 and BA.4/BA.5 compared to BA.1 or BA.2 (Hachmann *et al*., 2022; Quandt *et al*., 2022). A similar observation was made with BBIBP-CorV (Sinopharm) vaccinated individuals with and without Omicron PVI (Yao et al., 2022).

A challenge for informing vaccine strain selection with variant-specific antibody titers is the need to accommodate and interpret antibody breadth upon increasingly complex time-varying antigenic histories derived from infection, vaccination, or both (hybrid immunity). Compounding this challenge is the need to predict humoral immunity against yet-to-emerge variants. Antigenic cartography is a statistical method that geometrically interprets antibody titers, positioning variants in antigenic space based on how they are neutralized by primary exposure sera (Smith et al., 2004). Techniques that build on antigenic cartography, like antibody landscapes, evaluate how immunological breadth changes following re-exposure versus primary exposure and predict titers against parts of antigenic space that are not yet occupied by variants (Fonville et al., 2014). Very few antigenic maps have been made of SARS-CoV-2, likely because antigenic cartography requires well-characterized sera from individuals with primary exposure to distinct, sequence confirmed variants or experimentally inoculated animals (Amanat et al., 2021; Lusvarghi *et al*., 2022; Mykytyn et al., 2022; Neerukonda et al., 2021b; Rössler et al., 2022; van der Straten et al., 2022; Wilks et al., 2022). While the antigenic maps of SARS-CoV-2 published to date agree on the antigenic relationships between the ancestral strain, Delta, Beta, and Omicron, the positions of BA.2, BA.2.12, and BA.4/BA.5 remain uncertain.

In this study, we used antigenic cartography to measure the antigenic divergence among the major SARS-CoV-2 variants including BA.2, BA.2.12.1, and BA.4/BA.5 based on well-characterized sera from a longitudinal cohort following primary COVID-19 cases with sequence-confirmed variant infection histories. We complemented this with similar measurement from a separate cohort of uninfected individuals before and after their 2^nd^ and then 3^rd^ doses with mRNA vaccines. We then used antibody landscapes and other related tools to infer antigenic space and evaluate shifts in immunodominance following vaccination and infection. This approach enabled us to quantify the gain in magnitude and breadth following two or three doses with ancestral strain-based vaccines and Omicron PVI compared to boosting with the ancestral strain-based vaccine alone. Together, this analytical framework provides detailed information on breadth of observed and predicted immunity to SARS-CoV-2 variants following primary and subsequent exposure and informs selection of optimal vaccination strategies. This approach along with many other considerations, including variant surveillance, operational logistics, and availability of candidate vaccines and clinical data, can be used by public health authorities when making final recommendations for vaccine composition.

## RESULTS

### Neutralization of VOCs by primary infection antisera

To measure the antigenic relationships among variants using well-characterized primary infection antisera, we used sera from SARS-CoV-2 infected participants from the Epidemiology, Immunology, and Clinical Characteristics of Emerging Infectious Diseases with Pandemic Potential (EPICC) study (**Table S1**) (Epsi et al., 2022). We identified n=47 serum samples collected 8 to 51 days post symptom onset (mean=28 days) from individuals with natural primary infections with 21 distinct variants (all prior to vaccination). All these unvaccinated individuals had sequenced, genotyped infecting viruses, matched with clinical and demographic data (**Table S1 and S3**). An additional 31 convalescent serum samples with known infecting genotype were purchased from Boca Biolistics (Pompano Beach, FL, USA, **Table S4, Table S5**) (Neerukonda *et al*., 2021b). We also included an additional Beta-infected case from an unrelated FDA CBER study (see **Methods, Table S1**). Each of the 78 serum samples were titrated against a panel of SARS-CoV-2 lentiviral pseudoviruses representing the major variants, including variants of concern (n=15), including BA.1, BA.1.1, BA.2, BA.2.12.1, and BA.4/BA.5. A subset of sera was titrated against pseudoviruses consisting of ancestral strain with the D614G mutation (D614G) with one of seven individual point mutations introduced into the spike protein (**Table S6**).

Neutralization titers (ID50) for sera against each variant were grouped by infecting variant and shown in **Fig. 1**. For each serum group, significant differences in magnitude were observed across the variant panel, although the pattern of neutralization depends on the infecting variant. Variant that temporally preceded Omicron also generally have lower titers against Omicron variants. The highest geometric mean titer (GMT) across sera was generally to the infecting variant, and Alpha and Delta sera showed higher titers against the infecting variant compared to D614G. Among the pre-Omicron infections, and in agreement with previous data (Aleem et al., 2022; Collier et al., 2021; Guo et al., 2022; Ho et al., 2021; Mlcochova et al., 2021; Uriu et al., 2021; Wang et al., 2021a; Wang et al., 2021b; Wibmer et al., 2021), the titers of Alpha, Delta, Epsilon, and Lambda convalescent serum against Beta, Gamma and Mu variants were in general lower than against other pre-Omicron variants. Titers graphed according to emergence of the variants are shown in Fig. 1 panel K, and fold changes relative to the infecting variant are shown in Fig.1 panel L.

**Figure 1.**
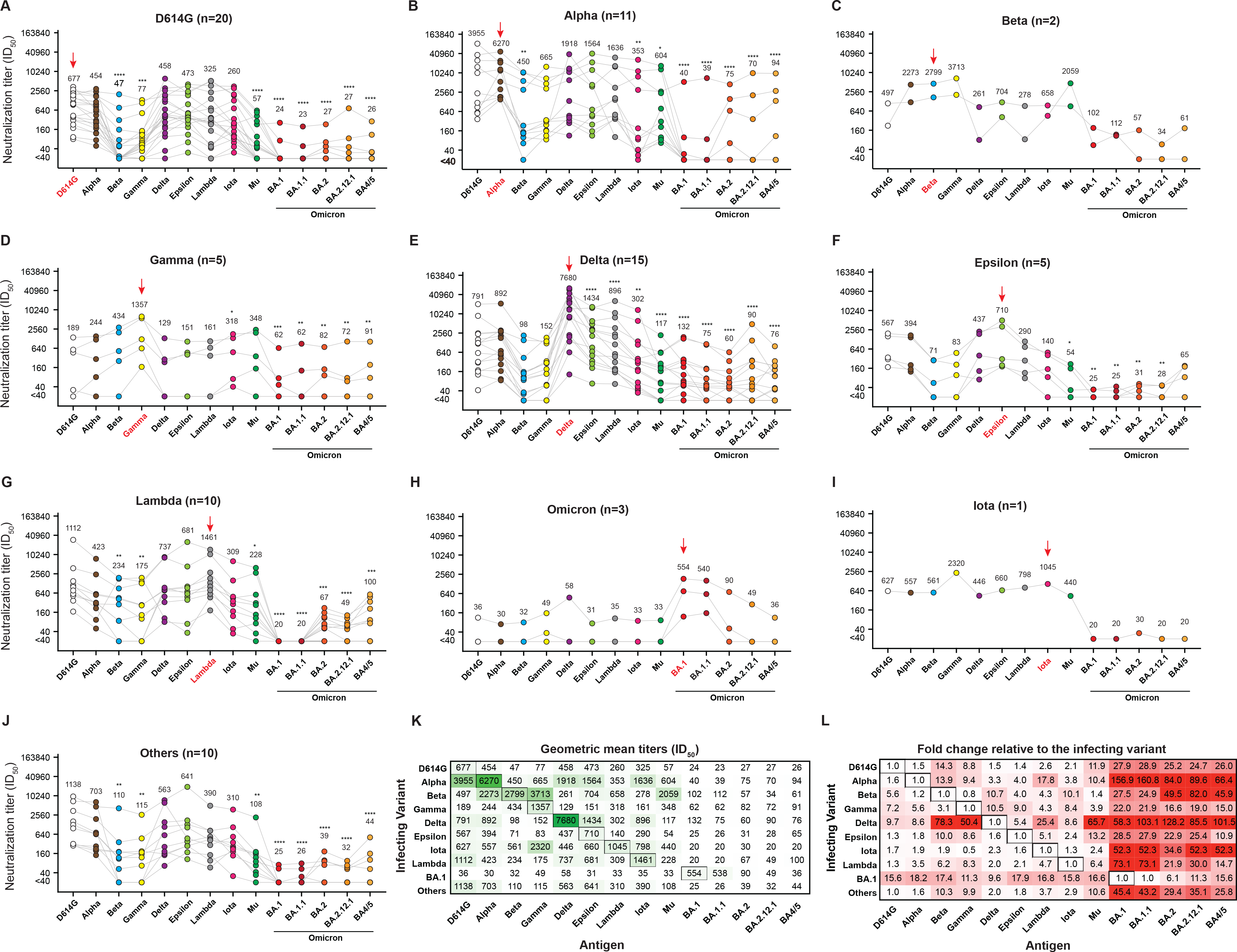
Neutralizing antibody titers (ID50 values) against SARS-CoV-2 variant pseudoviruses for primary infection convalescent sera from individuals infected by different Variants of Concern (VOCs). Sera from **A**) wildtype variant (D614G), **B**) Alpha, **C**) Beta, **D**) Gamma, **E**) Delta, **F**) Epsilon, **G**) Lambda, **H**) Omicron (BA.1 or BA.1.1), I) Iota, and **J**) other variants. Each grey line corresponds to one serum sample. Red arrow denotes the infecting variant. Geometric mean neutralizing antibody titers (GMT) are listed for each variant. Significance values for each variant are shown relative to the infecting variant. **K**) GMTs from panels A-J for sera from the infecting variants (rows) against all measured antigens (columns). Cells are shaded based on GMT, and serum-antigen pairs with larger titers have darker shades of green. **L**) Fold reduction in titer for each serum-antigen pair relative to the titer to the infecting variant (boxed in black). Each cell value represents the average fold change across all serum samples with the same exposure history, and darker red cells denote larger relative reductions in titer. For all neutralization assays, serum was diluted 1:40 followed by three-fold serial dilutions. Neutralization assays were performed twice, each with an intra-assay duplicate. Neutralization curves were fitted using nonlinear dose-response regression. Titers measuring below the lowest serum dilution of 1:40 were treated as 20 for statistical analysis. Statistical analysis was performed on the paired samples using the Friedman test, followed by post hoc Dunn’s multiple comparison tests. P values for comparisons between the groups are shown, where *P ≤ 0.05, **P ≤ 0.01, ***P ≤ 0.001, and ****P ≤ 0.0001.

### Primary infection antigenic maps show Omicron variants BA.2, BA.4/BA.5, and BA.212.1 as antigenically distinct from BA.1 and shifted toward the Beta variant

We used antigenic cartography to interpret all 1240 primary natural infection neutralizing antibody titer measurements and quantify the breadth of immunity across variants. Using a form of multi-dimensional scaling, each strain and serum is positioned in high dimensional Euclidean space such that the distance between points corresponds to the measured neutralizing antibody titer. The closer a serum (square) is to a variant (circle), the higher the titer for that serum to that antigen. Overall, we find that the sera cluster near their respective infecting variants, as expected. Using cross-validation, excluding 10% of titers for each test, we determined that two dimensions was sufficient to accurately fit the titers (average root mean squared error of 1.36 antigenic units, variance of 0.04); 3D maps are shown in **Fig. S1**. Points on the antigenic map were well coordinated and robust to measurement error in the assay as well as bootstrapping of individual viruses and sera (**Fig. S2**).

Consistent with previously published SARS-CoV-2 antigenic maps (Amanat *et al*., 2021; Lusvarghi *et al*., 2022; Mykytyn *et al*., 2022; Neerukonda *et al*., 2021b; Rössler *et al*., 2022; van der Straten *et al*., 2022; Wilks *et al*., 2022), we found four major variant groups that define the observed limits of SARS-CoV-2 antigenic space (**Fig. 2A**). These clusters generally correspond to groups with shared amino acid changes in the spike receptor binding domain, listed in parentheses below for each variant below. The variants clustered nearest to the ancestral strain (D614G) were Alpha (N501Y), Epsilon (L452R), and individual point mutations introduced into D614G (N501Y, L452R, T478K, R346K, and K417N). Lambda (L452Q and F490S) is only slightly further to the right and Delta slightly below the ancestral strain (L452R and T478K). To the top and right of the ancestral is the Beta cluster, consisting of Mu (E484K and N501Y), Gamma and Beta (E484K, N501Y, K417N/T), and D614G with both mutations E484K and N501Y. Iota (E484K), R.1 (E484K), and D614G with E484K are between the ancestral and Beta cluster, likely because they lack the additional antigenic mutation at N501Y. Omicron BA.1 and BA.1.1 are to the right and are most distant from the ancestral variant (123.3-fold difference). Both contain additional mutations in the receptor binding domain that are not observed in other VOCs, while BA.1.1 also contains R346K (**Fig 2E**). Strikingly, we found that BA.2.12.2, BA.2, and BA.4/BA.5 retain a large antigenic distance from the ancestral strain but are shifted away from BA.1 and BA.1.1 and toward the Beta cluster, supporting a recent observation that BA.2.12.1, BA.4 and BA.5 escape antibodies elicited by Omicron infection (Cao et al., 2022). BA.2 has numerous changes relative to BA.1 and BA.1.1 but is closely related to BA.2.12.1 and BA.4/BA.5 (**Fig. 2E**).

**Figure 2.**
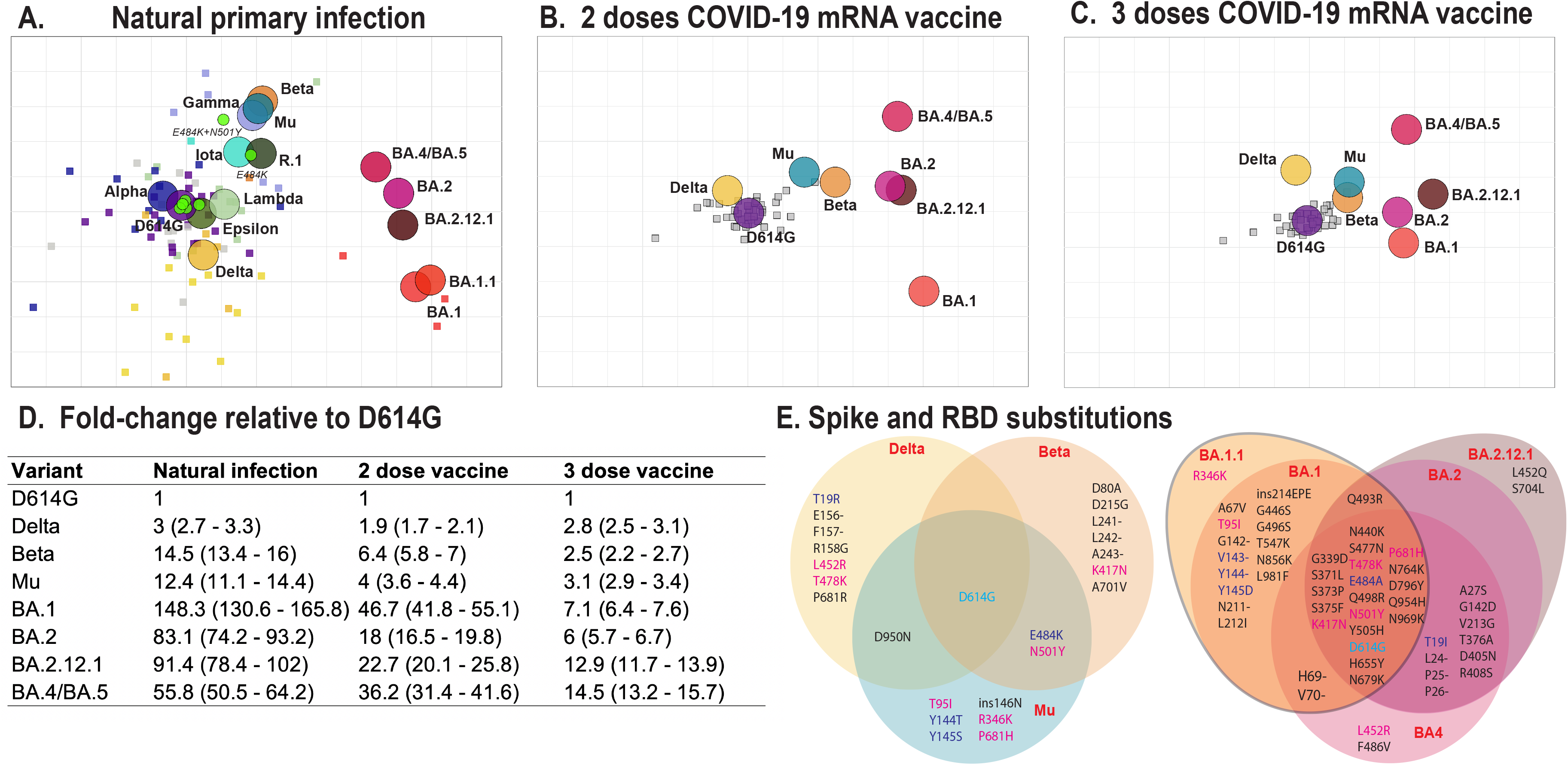
Antigenic maps made with neutralizing antibody titers from single-antigen exposure sera demonstrate BA.1, BA.2, BA.2.12.1, and BA.4/BA.5 are most antigenically distinct from other VOCs. Antigenic maps were made using antigenic cartography with titers for **A**) sera collected after convalescent primary infection with distinct VOCs and sera from uninfected individuals who received **B**) 2 doses or **C**) 3 doses of WT mRNA COVID-19 vaccines. Each grid-square side corresponds to a two-fold dilution in the pseudovirus neutralization assay. Antigenic distance is measured in any direction on the grid. Antigens are shown as circles and are labeled. Sera are shown as squares and are colored by infecting variant. **D**) Fold-difference in neutralization with 95% confidence intervals from the ancestral strain to each other variant on each map. For example, a fold-difference of four corresponds to two grid-squares on the antigenic map. **E**) Substitutions in the spike and receptor binding domains for all variants used in this study.

### Antigenic maps of two and three dose COVID-19 mRNA vaccine sera show distinct antigenic relationships for Omicron variants

We next measured the neutralization breadth of sera collected from n=39 health care workers after two and three doses with COVID-19 mRNA vaccines as part of the Prospective Assessment of SARS-CoV-2 Seroconversion (PASS) study (see **Methods, Table S2**). Sera were titrated against D614G, four Omicron variants, BA.1, BA.2, BA.2.12.1, and BA.4/BA.5, and Beta, Mu, and Delta. The last three were chosen because they represented the most distant clusters on the convalescent map in Fig. 2A. After two doses, titers were highest to the ancestral strain and lowest to BA.1. (**Fig. 3A**). Titers were low but slightly higher against BA.2, BA.2.12.1 and BA.4/BA.5. Both BA.1 and BA.4/BA.5 had the fewest titers above assay cut-off (<40). In contrast, the third vaccine dose significantly boosted GMTs to all variants (P<0.0001), with BA.2 having the highest titers among the Omicron-like viruses (831), followed by BA.1 (700), BA.2.12.1 (395), and BA.4 (355) (**Fig. 3B**).

**Figure 3.**
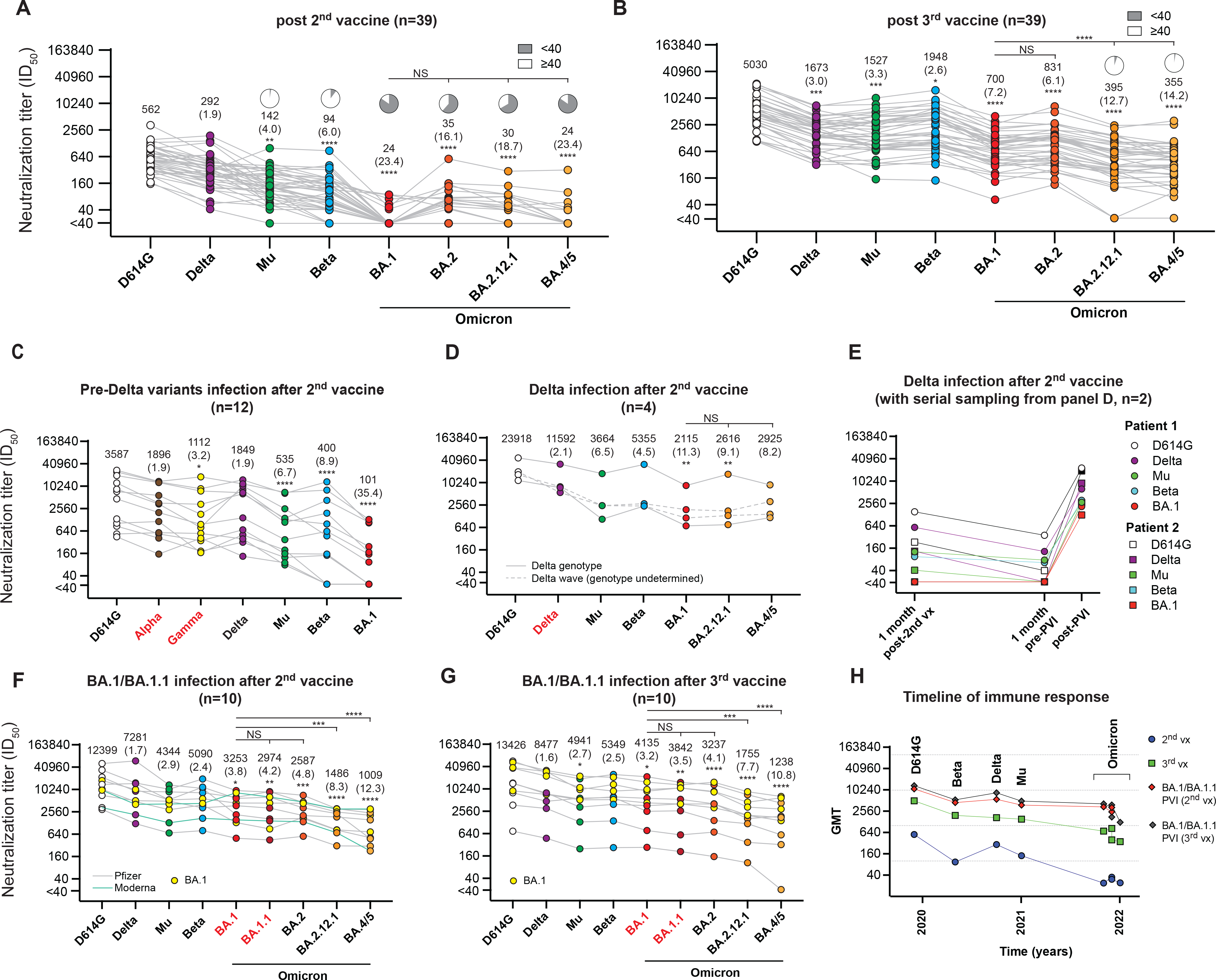
Neutralizing antibody titers (ID50 values) against variant pseudoviruses from post-vaccination sera with and without post-vaccination infection (PVI). Sera are from individuals who received **A**) 2 doses of a WT mRNA COVID vaccine or **B**) 3 doses of a WT mRNA COVID vaccine. Serum samples were obtained about 5-6 weeks following the last vaccine dose. Neutralizing antibody titers PVI after 2 doses of WT mRNA COVID vaccine in individuals infected with the **C**) pre-Delta wave (Alpha or Gamma or others), **D-E**) Delta, or **F-G**) Omicron (BA.1/BA.1.1). Each grey line corresponds to one serum sample. GMT are listed for each variant. Significance values for each antigen are shown relative to the titer against D614G. Two of the Delta wave PVI serum samples in panel D) were measured at multiple time points, shown in panel E), from 1-month post-vaccine dose 2, and 1 month before and after PVI. Panel F shows titers from individuals with an Omicron (BA.1/BA.1.1) PVI 2-10 months after the second vaccine, while Panel G shows titers from individuals with an Omicron (BA.1/BA.1.1) PVI 1-5 months after the third vaccine. **H**) The GMT of individual variant after vaccination with or without PVI by timeline. For all neutralization assays, serum was diluted 1:40 followed by three-fold serial dilutions. Neutralization assays were performed twice, each with an intra-assay duplicate. Neutralization curves were fitted using nonlinear dose-response regression. Titers measuring below the lowest serum dilution of 1:40 were treated as 20 for statistical analysis. Statistical analysis was performed on the paired samples using the Friedman test, followed by post hoc Dunn’s multiple comparison tests. P values for comparisons between the groups are shown, where *P ≤ 0.05, **P ≤ 0.01, ***P ≤ 0.001, and ****P ≤ 0.0001. NS: no significance; vx: vaccine. Pie charts indicate percent of serum sample above the lowest tested (1:40). Numbers in parentheses indicate fold reduction in titer relative to D614G.

Because antigenic maps can be made from sera with multiple exposures to the same antigen, we also used antigenic cartography to interpret neutralizing antibody titers for sera collected after two and three vaccine doses. Titers were accurately fit as antigenic maps in either one or two dimensions (**Fig. S1**) but coordination was less accurate than for the natural infection map (**Fig. S2**). We found that the antigenic relationships of variants on the two doses vaccine antigenic map were similar to the natural infection map (**Fig. 2A and B**), with Beta, Mu, and Delta closer to the ancestral strain and BA.1 furthest from the ancestral strain, followed by the other Omicron variants. In contrast, a marked change in immunodominance was observed when the same vaccinated individuals received their third dose (**Fig. 2C**). The antigenic distance between the ancestral strain and BA.1 and BA.2 reduced to 7.1- and 6-fold difference, while BA.2.12.1 and BA.4/BA.5 remained more divergent, at 12.7 and 14.4-fold difference (**Fig. 2D**). This observation suggests that the third dose specifically increased more breadth to BA.1 and BA.2 but less to BA.2.12.1 and BA.4/BA.5. A similar phenomenon was observed for Delta, which remained a similar antigenic distance following the third dose, compared to Beta and Mu, which shifted closer to the ancestral strain (Fig. 2C and 2D). These results suggest that booster vaccination with the ancestral variant selectively boosts antibodies to epitopes present on some variants but not others, in a way that is distinct from the antigenic distances measured based on primary infection serological responses.

### Two or three doses with COVID-19 mRNA vaccines followed by an Omicron PVI provided broader antibody breath than three vaccine doses alone

Although there are important differences between vaccination and infection, comparing the breadth of immunity between individuals with Omicron PVIs to those with only three vaccine doses may be considered a proxy for comparing the breadth induced by different boosting strategies. We measured neutralizing antibody titers for individuals with PVIs in the EPICC study and compared their responses to those with only two or three vaccine doses from the PASS study (**Table S1, Table S3**). In agree with previous report (Richardson et al., 2022), individuals with two vaccine doses followed by an Omicron (BA.1 or BA.1.1) PVI had high titers against previously circulating variants but lower against the early Omicron variants BA.1 and BA.2, and much lower for later variants BA.2.12.1 and BA.4/BA.5 (**Fig. 3F and G**). Among this group, some individuals with PVIs 8-10 months post-vaccination had a broader and higher magnitude response compared to individuals with PVIs 2-3 months post-vaccination. Individuals with three vaccine doses followed by Omicron (BA.1 or BA.1.1) PVI also had higher titers against BA.1, followed by BA.2, BA.2.12.1, and BA.4/BA.5, and overall higher titers than were observed after three vaccine doses alone. Individuals with two or three doses and then Delta infection had the highest titers against all VOCs (**Fig. 3D**). However, individuals with pre-Delta wave infection soon after two doses of vaccines had lower titers against all variants compared to other PVI groups, but higher titers than those in the two-dose vaccine group (**Fig. 3C**). Even if the neutralization titers dropped to background after the second vaccine, the PVI boosted high neutralization titers against all variants, indicating strong back-boosting to earlier variants (**Fig. 3D**).

We used antibody landscapes to evaluate breadth across antigenic space, assuming antigenic distance measured by primary infection antisera is related to antigenic relationships seen by repeat exposure antisera. The x and y dimension correspond to the original two-dimensional antigenic map made with primary natural infection antisera. In the third dimension, at each virus position on the map, the height of the landscape corresponds to measured neutralization titer for that serum against the virus. Here, we use the method used by Rössler (Rössler *et al*., 2022) and assume that antibody landscapes for individuals with multiple prior exposures cones with slopes that can deviate from 1. We first tested whether infection with an antigenically distinct variant, such as BA.1 in those with prior vaccination induced broader immunity than a third dose with the ancestral strain. We found that individuals with two vaccine doses followed by Omicron (BA.1/BA.1.1) PVI had a more gradual slope, indicating broader immunity, than landscapes for individuals who received three doses of ancestral vaccine (**Fig. 4A and B**). Further, we found that Omicron (BA.1/BA.1.1) PVIs in those with 3 prior vaccine doses also broadened immunity beyond what was induced by the third vaccine dose, with a less steep slope and higher magnitude titers, consistent with a stronger, broader response (**Fig. 4B and C**). Notably, individuals with two or three doses and Omicron PVIs had broader immunity against Delta and Omicron variants BA.1, BA1.1, BA.2, BA.212.1, and BA.4/BA.5. Further, we found no substantial difference in breadth or magnitude between those with two versus three doses prior to Omicron PVI either from the raw titer data (**Fig. 3F, G, and H**) or antibody landscapes (**Fig. 4B and C**). Thus, an additional boost with the ancestral-based vaccine did not provide benefit in antibody responses..

**Figure 4.**
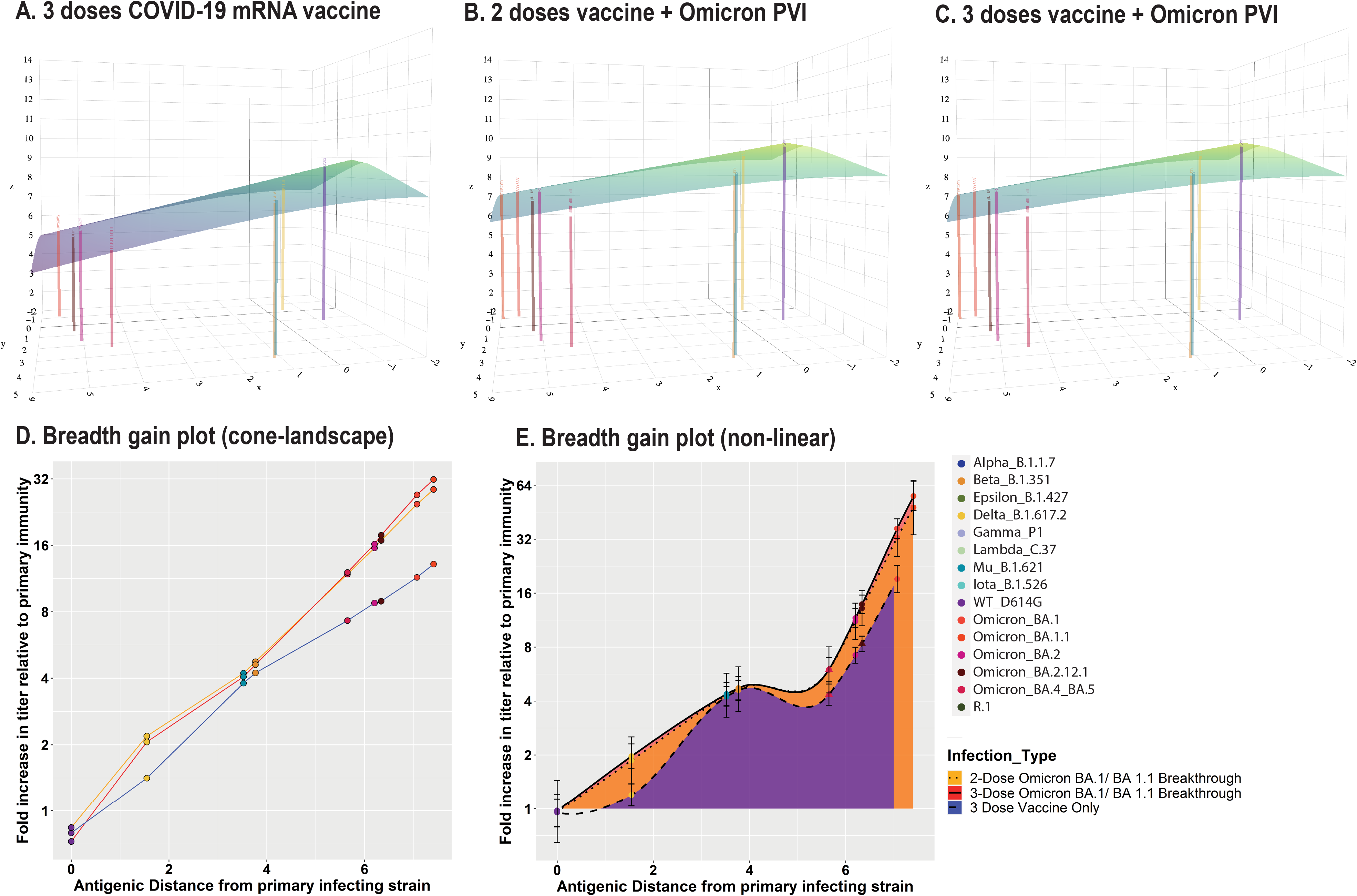
Antibody landscapes and breadth gain plots show that individuals with PVIs have a large gain in both breadth and magnitude compared to those with three mRNA vaccine doses alone. Antibody landscapes are shown for individuals with **A**) 3 doses of mRNA vaccine **B**) 2 doses of mRNA vaccine followed by Omicron PVI, and **C**) 3 doses of mRNA vaccine followed by Omicron PVI. The x and y-axis on each landscape correspond to the 2D antigenic map constructed from convalescent sera in Fig. 2A, with colored points representing the locations of each measured antigen. The z-axis in each landscape represents the interpolated log GMT for all individuals with that exposure history against each antigen. The average landscape for each serum group was constructed by fitting landscapes for each individual serum sample assuming that all landscapes with the same infection history have the same slope, with peak equal to the maximum observed titer value against any one of the measured antigens. The location of the peak titer value was fitted separately for each individual and then subsequently averaged. The colored lines represent the expected average log GMT for individuals with a particular infection history against each measured antigen. The color of the landscape, like the z- axis, corresponds to estimated log GMT across antigenic space. **D**) Breadth gain plots of the antibody landscapes in A-C for vaccinated individuals who received either a third mRNA vaccine dose, an Omicron PVI, or both. The x-axis represents the antigenic distance from the primary convalescent sera antigenic map (Fig. 2A) between the primary exposure variant and each measured antigen. Each unit on the y-axis represents the log-fold increase in titer against a particular measured antigen beyond a primary infection response and is used to compare the relative breadth of different exposure histories. Conceptually, the x-axis represents the antibody landscape for primary exposure sera projected into 1-dimension. The y-axis is in units of antigenic distance, corresponding to log fold-difference in neutralizing antibody titers (the same units as the antigenic map). **E**) Same as D, showing gain values for each set of sera with the same infection history interpolated from a loess fit (non-linear). Error bars represent the mean and 95% confidence intervals for at each measured antigen. Shading colors and lines denote the type of infecting variant and the number of vaccine doses received.

### Detailed characterization of antibody breadth reveals boosting of immunity is lower to some variants, including BA.2.12.1 and BA.4/BA.5

When we examined the residuals of our antibody landscapes, we observed lower titers against BA.2.12.1 and BA.4/BA.5 than predicted by the landscape. We developed a new method, which we call the ‘breadth gain’ plot, to compare linear (cone-shaped landscape) and non-linear increases in antigenic breadth relative to the primary antigenic map (**Fig. 4D, Tables S7 to S10**). We use this method to quantify the extent to which Omicron PVIs provided broader immunity than three doses with the ancestral vaccines. The breadth gain plots for the cone-landscapes are shown in **Fig. 4D**. The non-linear breadth gain plots (**Fig. 4E**) Omicron PVIs following two or three doses of vaccination provided significantly broader protection than three vaccine doses against Omicron variants BA.1, BA.1.1, and BA.2, as well as Delta. Compared to three vaccine doses alone, three vaccine doses and Omicron PVI induced significantly greater breadth to BA.4/BA.5, while this difference was not significant for those with two doses with Omicron PVI. Interestingly, however, boosting was more limited to BA.2.12.1 and BA.4/BA.5 as compared to BA.1, both following three vaccine doses or vaccination with Omicron PVI. Together, these observations suggested there may be ‘valleys’ in the antibody landscape, indicating regions of antigenic space with lower-than-expected titers. This could occur if certain epitopes present in distinct variants are preferentially boosted.

## DISCUSSION

When considering a vaccine antigen to optimize protection, breadth should be framed in the context of circulating variants, as well as an antibody titer that would be needed for protection. Characterizing antibody breadth is complex, especially as SARS-CoV-2 continues to evolve and host exposure histories become more varied with each new wave of variants and vaccination campaigns. In this study, we provide an analytic framework to account for these antigenic determinants of humoral immunity. We used sera from individuals following primary infection, vaccination, and PVIs to examine how the antigenic characteristics of SARS-CoV-2 have diversified over time. We also introduced methods for quantifying immune breadth and magnitude that capture antigenic complexities when evaluating both primary and booster vaccination strategies. This approach can be incorporated into a larger toolkit for identifying vaccine strategies that broadly boost across variants.

We cannot know how SARS-CoV-2 will evolve as it adapts to the human population with rapidly changing background immunity, so judgements will have to be made using best available data. Antigenic cartography provides a useful framework for evaluating the breadth of neutralizing responses across variants and has been used to monitor the antigenic evolution of influenza (Russell et al., 2008) and dengue viruses (Katzelnick et al., 2021), among other pathogens. Our antigenic maps of pre-Omicron variants agree with previously published antigenic maps of SARS-CoV-2 (Amanat *et al*., 2021; Lusvarghi *et al*., 2022; Mykytyn *et al*., 2022; Neerukonda *et al*., 2021b; Rössler *et al*., 2022; van der Straten *et al*., 2022; Wilks *et al*., 2022), likely due in part to use of similar pseudovirus assays across laboratories. We find strong clustering of the original variants with shared amino acid positions, indicating that specific amino acid changes determine antigenic phenotype. The position of BA.2 on our map agrees with an experimental animal antigenic map (Mykytyn *et al*., 2022) but differs from a map made with natural infection antisera (Rössler *et al*., 2022) . We further showed that BA.2.12.1 and BA.4/BA.5 are closer to the Beta cluster, which may provide insight for related epitopes in these variants.

Our findings demonstrate differences in immunodominance hierarchies among variants, which could have implications for selection of new vaccine antigens. We found that a third vaccination with the ancestral variant selectively boosts antibodies to epitopes present on some variants but not others; specifically, breadth to BA.1 and BA.2, but not to BA.2.12.1 and BA.4/BA.5, was increased. A similar phenomenon was observed for Delta, which remained a similar antigenic distance following the third dose, compared to Beta and Mu, which shifted closer to the ancestral strain. A previous study found that additional mutations in BA.2.12.1 (L452Q), and BA.4/BA.5 (L452R, F486V and the deletion in 69-70) help explain antigenic differences relative to BA.2 for three dose vaccinee sera (Wang *et al*., 2022). However, to our knowledge, the specific mutations that explain why antigenic distances measured based on sera from primary infection and two vaccine doses differ from those with three vaccine doses remains to be explained. If antigenic distance can be used to select an antigen for inducing breadth, then BA.4 might be expected to induce broader immunity than BA.1 in highly vaccinated populations. Alternatively, if it is beneficial to boost with variants in related regions of antigenic space, vaccination with Beta/Mu/Gamma-type variant might provide broader protection against BA.4/BA.5-type strains than either ancestral vaccination or BA.1/bivalent boosting because BA.4/BA.5 is shifted toward the Beta cluster (Launay et al., 2022).

We measured both the breadth and magnitude of antibody titers for individuals with multiple prior exposures using antibody landscapes. Antibody landscapes can be constructed using various methods depending on the amount of data, ranging from a simple plane (Wilks *et al*., 2022) to fitting a continuous surface with multiple datapoints (Fonville *et al*., 2014). Given the more limited number of variants available for SARS-CoV-2, we fit antibody landscapes assuming they are shaped like a cone, with a variable peak location and slope (Rössler *et al*., 2022). This builds on an intrinsic feature of Euclidean antigenic space, which is that an antibody landscape for a primary infection serum that is perfectly fit by the antigenic map is a cone with a slope of 1, such that each two-fold drop in distance in the z-axis corresponding to a 2-fold drop in the x and y-dimensions. Using this approach, we found that repeated exposure to the ancestral strain from an mRNA vaccine broadened immunity to previously circulating strains after the third dose, indicating strong back-boosting, as well as to the Omicron variants. However, individuals with two vaccine doses followed by Omicron PVIs had even flatter landscapes and hence broader immunity against ancestral and Omicron-lineage variants relative to individuals vaccinated with only the ancestral variant. We also found that three doses with Omicron PVI induced broader responses than three vaccine doses alone, suggesting that individuals who already received a third dose with the ancestral strain could still potentially broaden their immunity by receiving an Omicron vaccine. Finally, we evaluated whether individuals with a third vaccine dose followed by Omicron PVI obtained an extra gain in breadth due to their additional vaccine dose, but found their responses were similar to individuals with only two doses and Omicron PVI, at least in the short-term following vaccination. Notably, both those with two and three vaccine doses and Omicron PVI had greater breadth across a range of variants, including Delta. Delta is in the lower, more distant part of the antigenic map, suggesting back-boosting of responses to earlier variants by Omicron PVI. The effect may provide protection against possible future antigens that could occupy that region of space.

To test for hills and valleys in antibody landscapes indicating preferential boosting of certain variants and weaker-than-expected boosting against other variants, we developed an analysis called the breadth gain plot. These analyses indeed showed lower responses to BA.2.12.1 and BA.4/BA.5 in all vaccinated and PVI groups than expected based on measured antigenic distances. This feature was not captured by our cone-based landscapes, which assume that because BA.21.2.1 and BA.4/BA.5 are closer to the ancestral strain than BA.1 on the antigenic map, sera with high titers to both the ancestral strain and BA.1 would also have high titers to BA.2.12.1 and BA.4/BA.5. However, this finding matches our antigenic analyses showing that three vaccine doses preferentially boosted BA.1 and BA.2 but not BA.2.12.1 or BA.4/BA.5. Collectively, these results point to a complex immunodominance pattern in which responses are boosted (whether by repeated ancestral vaccination or Omicron PVI) against epitopes that present to a lesser degree on BA2.12.1 and BA.4/BA.5 compared to BA.1. The ‘valley’ in the landscape for BA.4/BA.5 is notable and may indicate the type of immunodominance patterns previously observed in viruses that have circulated in human populations for decades, such as influenza. For example, during the 2013-2014 flu season, the H1N1 virus infected large numbers of middle-aged adults (Linderman et al., 2014). Subsequent analyses showed that several mutations on the virus, occurred at epitopes that were targeted by antibody responses in middle-aged adults, likely due to their prior exposure history. Similarly, vaccine responses may also be shaped by immune imprinting from prior vaccines or different SARS-CoV-2 variant exposures (Reynolds et al., 2022). Thus, while infection and or vaccination with BA.1 could increase immunity to current Omicron variants, whether use of the BA.1 as a first Omicron antigenic exposure could affect subsequent immunodominance patterns to the other Omicron or future variants remains to be studied.

Overall, our results show how the antigenic co-evolution of the SARS-CoV-2 and its immune response among the host human population become more elaborate with time. We present methods that can be used to characterize the breadth of immune responses to COVID-19 that account for diverse antigenic exposures. We show that Omicron PVIs generally induces broader immunity than boosting with the ancestral vaccine, and that additional exposure to both ancestral and BA.1 antigens can even increase breadth against BA.4/BA.5. However, the breadth gained to BA.4/BA.5 is lower than against BA.1, even though BA.4 is antigenically closer to the ancestral strain, indicating complex immunodominance patterns. Understanding the mechanism behind these immunodominance shifts and carefully quantifying immune breadth may become increasingly important for developing vaccination strategies against future COVID-19 strains. Further, scientific insights into the evolution of antigenic diversity for SARS-COV-2 could also shed light on older antigenically complex diseases by illustrating how the immunodominance patterns and complex landscapes that we currently observe may have evolved.

## LIMITATIONS OF THE STUDY

Our study used convalescent serum samples that were collected at different times point post COVID-19 diagnosis (3-51 days), which could have affected measured magnitude and breadth. Some of the commercial serum samples reported sample collection dates soon after symptom onset. We only used samples that had high neutralization titers, suggesting that the infection may have been well underway before reported symptom onset. To maximize serum coverage of VOCs on the antigenic map, we included 10 samples that were not fully genotyped but assigned variant infections based on dates of circulating variants at the time of sample collection. Ideally antibody landscapes are constructed by fitting interpolated surfaces across antigenic space as in Fonville 2014, but there are not yet enough distinct VOCs for this method. The emergence of future variants, titration with additional subvariants, or generation of mutant pseudoviruses that probe unoccupied areas of antigenic space may make more comprehensive antibody landscape analyses possible. There were too few individuals with PVIs with other variants to evaluate statistical significance with landscapes and breadth gain plots. Future studies on larger numbers of individuals or samples from clinical trials will provide further information on how sequential exposure to distinct antigens covers antigenic space. Finally, while neutralizing antibody titers measured with pseudovirus neutralization assays are correlated with protection, our study does not directly provide information on protection against VOCs. Further studies incorporating antibody landscapes with disease outcome data will provide further insights into how immune breadth across antigenic space is associated with clinical protection.

## STAR METHODS

### Data and materials availability

All data and code associated with this study are in the paper or supplementary materials. Sera samples are subject to an MTA and sera availability.

### Experimental Model and Subject Details

#### Ethics statement

The PASS (Protocol IDCRP-126) and EPICC (Protocol IDCRP-085) studies were approved by the Uniformed Services University of the Health Sciences Institutional Review Board (IRB) in compliance with all applicable Federal regulations governing the protection of human participants. All PASS and EPICC study participants provided informed consent. The convalescent Beta sera, obtained from a traveler who had moderate-severe COVID-19 in the Republic of South Africa during the peak of the Beta (B.1.351) wave in January 2021, was obtained with informed consent and covered under the US Food and Drug Administration IRB approved expedited protocol # 2021-CBER-045.

#### Collection of sera from vaccinees with no history of infection: study population, setting and procedures

Details of the Prospective Assessment of SARS-CoV-2 Seroconversion (PASS) study protocol, including details of the inclusion/exclusion criteria, have been previously published (Jackson-Thompson et al., 2021). Inclusion criteria included being generally healthy, ≥ 18 years old, and employed at the Walter Reed National Military Medical Center (WRNMMC), Bethesda as a healthcare worker. Exclusion criteria included history of COVID-19, IgG seropositivity for SARS-CoV-2 (as determined by a binding antibody assay) and being severely immunocompromised at time of screening. The study was initiated in August 2020, with rolling enrollment and monthly research clinic visits to obtain serum for longitudinal SARS-CoV-2 antibody testing.

The subset of PASS uninfected vaccinee participants selected for analysis of sero-responses were those who received two doses of Pfizer/BNT162b2 vaccine by January 26, 2021, had no serological or PCR evidence of SARS-CoV-2 infection prior to two doses of vaccine, and had received a 3rd dose of Pfizer/BNT162b2 vaccine by Nov 18, 2021. No subject included in this sub-analysis of vaccinated participants had a clinically apparent PCR-confirmed SARS-CoV-2 infection during follow-up before sera collection. Participants’ serum samples were collected monthly through September of 2021, and then quarterly.

For the antibody binding assay used for screening at enrollment, serum samples were diluted 1:400 and 1:8000 and screened for immunoglobulin G (IgG) reactivity with SARS-CoV-2 spike protein and nucleocapsid protein (N), and four human coronavirus (HCoV) spike proteins using a multiplex microsphere-based immunoassay, as previously described (Laing et al., 2021).

#### Collection of post-infection convalescent sera in vaccinated and unvaccinated study participants: setting and procedures

The Epidemiology, Immunology, and Clinical Characteristics of Emerging Infectious Diseases with Pandemic Potential (EPICC) study is a cohort study of U.S. Military Health System (MHS) beneficiaries that includes enrollment and longitudinal follow up of those with a history of SARS-CoV-2 infection (Richard et al., 2021). Eligibility criteria for enrollment included presenting to clinical care with COVID-19-like illness and being tested for SARS-CoV-2 by polymerase chain reaction (PCR) assay. The EPICC study enrolled between March 2020 and April 2022. For this analysis derived from SARS-CoV-2 infections, EPICC enrollment occurred at eight Military Treatment Facilities (MTFs): Brooke Army Medical Center, Fort Belvoir Community Hospital, Madigan Army Medical Center, Naval Medical Center Portsmouth, Naval Medical Center San Diego, Tripler Army Medical Center, Walter Reed National Military Medical Center, and the William Beaumont Army Medical Center.

Study procedures for these participants with SARS-CoV-2 infection included collection of demographic data, and completion of a clinical case report form (CRF) to characterize the acute SARS-CoV-2 infection. Biospecimen collection included serial serum samples for immune response analysis and upper respiratory specimen swabs for genotyping of SARS-CoV-2. For all enrolled participants, we also abstracted MHS-wide healthcare encounter data from the Military Health System Data Repository (MDR) to determine comorbidities. Vaccination status was ascertained by the MDR record, the CRF and questionnaire self-report.

In addition to convalescent sera from EPICC participants, we included convalescent sera from two PASS participants with SARS-CoV-2 infection in August 2021 (during the Delta epidemic). Both participants were vaccinated with two doses of mRNA COVID-19 vaccine at the time of SARS-CoV-2 infection (**Table S1**).

#### Commercial convalescent sera

Convalescent sera from SARS-COV-2 infected donors were purchased from Boca Biolistics (Pompano Beach, FL). Samples were selected from the SARS-CoV-2 sequence inventory. Details about the serum donors are listed in **Table S4**. Genotypes of the infecting viruses are listed in **Table S5**.

#### Diagnosis of SARS-CoV-2 infection and genotyping of infections used for convalescent sera

For EPICC participants, SARS-CoV-2 infection was determined by positive PCR clinical laboratory test performed at the enrolling clinical MTF site, or a follow-up upper respiratory swab collected as part of the EPICC study procedures. The specific PCR assay used at the MTF varied. The SARS-CoV-2 (2019-nCoV) CDC qPCR Probe Assay research-use-only kits (Integrated DNA Technologies, IDT, Coralville, IA) was used as the follow-up PCR assay (used for specimens collected as part of the EPICC study). This CDC qPCR assay uses two targets of the SARS-CoV-2 nucleocapsid (N) gene (N1 and N2), with an additional human RNase P gene (RP) control. We considered a positive SARS-CoV-2 infection as positive based on a cycle threshold value of less than 40 for both N1/N2 gene targets.

Whole viral genome sequencing was performed on extracted SARS-CoV-2 RNA from PCR positive specimens using a 1200bp amplicon tiling strategy (https://doi.org/10.1093/biomethods/bpaa014). Amplified product was prepared for sequencing using NexteraXT library kits (Illumina Inc., San Diego, CA) and libraries were run on the Illumina NextSeq 550 sequencing platform. Genome assembly used BBMap v. 38.86 and iVar v. 1.2.2 tools. The Pango classification tool (version 4.0.6) was used for lineage classification (https://doi.org/10.21105/joss.03773). In a small minority of SARS-CoV-2 infections (**Table S3**), a Pangolin lineage was unable to be ascertained and either a Nextclade clade (https://doi.org/10.21105/joss.03773) was used and/or a Pangolin lineage was inferred by manual inspection of key lineage-defining amino acid substitutions. Dates of infection were also used as supplementary information to ascertain infecting genotype in such instances where spike sequence quality was lower.

In addition, the infecting genotype for one EPICC participant (Cov-83) **(Table S3**) was determined using the Illumina Miseq platform; cDNA synthesis was performed using the Superscript IV first-strand synthesis system (Life Technologies/Invitrogen, Carlsbad, CA). The ARTIC v3 primer set was used for multiplex PCR to amplify overlapping regions of the SARS- CoV-2 reference genome (MN908947.3). Primer and genomic alignment position information is available here: http://github.com/artic-network/artic-ncov2019/tree/master/primer_schemes/nCoV-2019/V1. The MinElute PCR purification kit (QIAgen, Valencia, CA) was used to purify PCR products and libraries were prepared with the SMARTer PrepX DNA Library Kit (Takara Bio, Mountain View, CA), with use of the Apollo library prep system (Takara Bio, Mountain View, CA). The quality of these libraries was evaluated using the Agilent 2200 TapeStation (Agilent, Santa Clara, CA); after quantification by real-time PCR using the KAPA SYBR FAST qPCR Kit (Roche, Pleasanton, CA), libraries were diluted to 10 nM.

Additionally, we included 10 convalescent sera with infecting genotype inferred by date of collection. Convalescent sera from seven vaccinated EPICC participants diagnosed with COVID-19 between 2/9/2021 and 4/2/2021 did not have corresponding viral sequence data to confirm the infecting genotype and were categorized as presumptive “pre-Delta” infections (**Fig 3C, Table S2**). The infecting genotype of two PASS participants with vaccine breakthrough infections were inferred by date of infection (late August 2021, annotated as presumptive Delta infections, **Fig 3D and 3E, Table S1**). Additionally, we included in all analyses sera from a traveler who had COVID-19 in the Republic of South Africa during the peak of the Beta (B.1.351) wave in January 2021 (collected under a separate CBER protocol, 2021-CBER-045) (**Table S1**) and this was annotated as a presumptive Beta infection.

### Method Details

#### Plasmids and Cell Lines

Codon-optimized, full-length open reading frames of the spike genes of SARS-CoV-2 variants in the study (Table S6) were synthesized into pVRC8400 or pcDNA3.1(+) by GenScript (Piscataway, NJ, USA).. The HIV gag/pol packaging (pCMVΔR8.2) and firefly luciferase encoding transfer vector (pHR’CMV-Luc) plasmids (Naldini et al., 1996; Zufferey et al., 1997) were obtained from the Vaccine Research Center (National Institutes of Health, Bethesda, MD, USA). 293T-ACE2-TMPRSS2 cells stably expressing human angiotensin-converting enzyme 2 (ACE2) and transmembrane serine protease 2 (TMPRSS2) (BEI Resources, Manassas, VA, USA; Cat no: NR-55293) (Neerukonda et al., 2021a) were maintained at 37°C in Dulbecco’s modified eagle medium (DMEM) supplemented with high glucose, L-glutamine, minimal essential media (MEM) non-essential amino acids, penicillin/streptomycin, HEPES, and 10% fetal bovine serum (FBS).

#### SARS-CoV-2 Pseudovirus Production and Neutralization Assay

HIV-based lentiviral pseudoviruses with desired SARS-CoV-2 spike proteins were generated as previously described (Neerukonda *et al*., 2021a). Pseudoviruses comprising the spike glycoprotein and a firefly luciferase (FLuc) reporter gene packaged within HIV capsid were produced in 293T cells by co-transfection of 5 µg of pCMVΔR8.2, 5 µg of pHR’CMVLuc and 0.5 µg of pVRC8400 or 4 µg of pcDNA3.1(+) encoding a codon-optimized spike gene. Pseudovirus supernatants were collected approximately 48 h post transfection, filtered through a 0.45 µm low protein binding filter, and stored at -80°C.

Neutralization assays were performed using 293T-ACE2-TMPRSS2 cells in 96-well plates as previously described (Neerukonda *et al*., 2021a). Pseudoviruses with titers of approximately 10^6^ relative luminescence units per milliliter (RLU/mL) of luciferase activity were incubated with serially diluted sera for two hours at 37°C prior to inoculation onto the plates that were pre-seeded one day earlier with 3.0 × 10^4^ cells/well. Pseudovirus infectivity was determined 48 h post inoculation for luciferase activity by luciferase assay reagent (Promega) according to the manufacturer’s instructions. The inverse of the sera dilutions causing a 50% reduction of RLU compared to control was reported as the neutralization titer (ID50). Titers were calculated using a nonlinear regression curve fit (GraphPad Prism Software Inc., La Jolla, CA, USA). The mean titer from at least two independent experiments each with intra-assay duplicates was reported as the final titer.

### Quantification and Statistical Analysis

#### Statistical analysis of neutralizing antibody titer data

One-way analysis of variance (ANOVA) with Dunnett’s multiple comparisons tests (variants compared to D614G-, variants compared to BA.1), two-way ANOVA for the comparison of different groups (i.e., two-dose vaccine vs three-dose vaccine) and geometric mean titers (GMT) with 95% confidence intervals were performed using GraphPad Prism software. The P values of less than 0.05 were considered statistically significant. All neutralization titers were log_2_ transformed for analyses.

#### Antigenic cartography

We used the Racmacs package (https://acorg.github.io/Racmacs/) for antigenic cartography analyses (Wilks *et al*., 2022). Antigenic maps are quantitative visualizations that fit antibody titers as Euclidean distances between primary infection antisera and variants. Datasets with diverse variants and primary infection sera to each variant are best for making meaningful geometric interpretations, i.e., antigenic maps. Racmacs implements a modified multi- dimensional scaling approach as previously described (Smith *et al*., 2004). Briefly, the virus best neutralized by each serum *j*, is defined as *bj*. *Nij* is the neutralization titer for serum *j* against virus *i*. The antigenic distance, *Dij*, for serum *j* to each virus *i* is defined relative to *bj*: *Dij* =log_2_(*bj*)- log_2_(*Nij*). The map Euclidean distance *dij* for each virus and serum is that which best fits the measured antigenic distance *Dij* in each number of dimensions. The optimal set of map coordinates for each serum and virus is identified by minimizing the stress function E=∑ije(*Dij*, *dij*) thousands of times from random starting coordinates using a conjugate gradient optimization. For titers measured above the assay lower limit of quantitation, the stress function minimized is (*D_ij_* - *d_ij_*)^2^. For titers below the assay lower limit of quantitation, the stress function minimized is 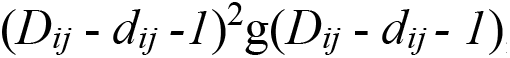, where 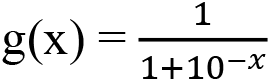. A unit of antigenic distance is equivalent to a two-fold dilution in neutralizing antibody titers. We performed various quality assessments for antigenic maps including evaluation of optimal dimensionality using cross validation, characterization of titer residuals, confidence coordination on the map, robustness to assay error and outlier viruses and sera.

### Antibody landscapes

We generated average landscapes for all serum samples with same overall infection history (e.g. 2 doses of ancestral vaccine, 2 doses of ancestral + Omicron PVI etc), following the general approach of Roessler and Netzel et al. (Rössler *et al*., 2022). We fitted three parameters for each landscape-the slope and x and y coordinates of the landscape peak. Let *x_p_* and *y_p_* represent the x and y coordinates of the landscape peak, and *s_k_* the slope. We assume that each of these parameters has the same value across all serum samples within the serum group. Let *C_j_* represent the column basis titer for each serum sample *j*. We assume that for each serum sample, the height of the landscape at the peak is equal to column base titer of the serum. Let *A_ip_* represent the antigenic distance between the peak and a particular measured antigen *i*. The predicted titer against measured antigen *i* for serum sample *j* is given by:

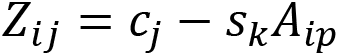

Let *T_ij_* denote the observed titer for serum sample *j* against measured antigen *i*. We use the function optim() in the R package stats to minimize the square error *E*, which is the sum of the difference between observed and predicted titers across all measured antigens and serum samples within the serum group:

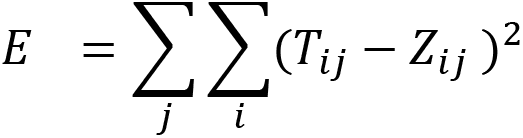

### Breadth Gain Plots

To obtain our measure for the relative breadth of the immune response for each individual (i.e., serum sample), each individual’s titer measurements against all measured VOCs were subtracted from the highest log titer for that serum across all antigens in the panel. This is the same type of titer normalization performed when constructing the antigenic map, and thus the units represent the same units of antigenic distance (i.e., 1 unit of distance corresponds to a 2-fold dilution). This is the “table distance” between the secondary infection serum and each measured antigen.

We then constructed our expectation of what this distance would be if the secondary immune response was identical to the primary infection response (i.e., if the secondary exposure did not change the response in any way). If the responses were identical, we would expect the peak of the landscape to be located on the antigenic map at the measured antigen closest to the primary infecting strain and decay uniformly from that point, with titers against other antigens decreasing proportionally to their map distance from the infecting strain.

In general, we expect that titers will decrease as antigenic distance between a serum and measured antigens increases, just as we observe for primary infection sera. However, secondary infection sera, unlike primary infection sera, were not used to generate the primary infection antigenic map and thus we do not have readily available serum coordinates. Instead, for sera whose infecting strain was one of the antigens that were measured (and that thus have coordinates on the antigenic map), we use those coordinates in lieu of serum coordinates and then calculate the distance on the antigenic map between that strain and all other strains against which the serum was titrated. Visually, this is analogous to drawing a series of concentric circles on the antigenic map centered on the infecting strain and radiating out to all other measured antigens on the map, and then calculating the radius of each circle. We refer to these distances as the “map distances” hereafter.

Not all infecting strains have corresponding antigens in the panel used for titration. For strains that were not included in the panel but were structurally similar to other panel strains, we used the panel strain that most closely resembled the infecting strain as the reference strain. A table of infecting strains and corresponding reference strains is in (Atmar et al.).

We arranged the measured antigens on the x-axis in order of increasing map distance (i.e., antigenic distance from the primary infecting strain). We then performed a Loess fit of the table distance between each measured antigen and the secondary infection serum, pooling all serum with the same secondary infection history category. The mean and standard error for each Loess fit was calculated at each measured antigen and was used to obtain 95% confidence intervals (mean +/- 2 times the standard error). When performing the Loess fit, we also interpolated the table distance a vector of 80 evenly spaced points between 0.1 and 8 antigenic units from the primary infecting strain to create a uniform curve that could be compared across different infection histories.

To obtain the deviance in immune breadth for each point in the deviation breadth plot, we subtracted map distance from the calculated table distance. Positive deviation values denote broader immunity than expected against a particular measured antigen, while negative values denote narrower immunity than expected. Loess fits were conducted using the loess() function in the R package stats with the default span setting of 0.75, and antigenic map coordinates were extracted using the R package racmacs. All analyses for the breadth plots was performed using the statistical software R version 4.2.0 (R Core Team, 2022).

For each measured antigen, we compared the difference in the fold breadth gain between groups of serum samples with different exposure histories. One-sided Mann Whitney tests were conducted for each pair of exposure groups, with the null hypothesis being that both distributions were the same, and the alternate hypothesis being that the broader exposure history’s distribution was greater than the distribution of the narrow exposure history. Classifications of “broad” and “narrow” exposure histories for purposes of one-sided comparison were made based on the breadth gain plot (**Figure 4E**). Tests were computed using the wilcox.test function in the R package stats.

## SUPPLEMENTAL INFORMATION

**Fig. S1. Evaluation of goodness of fit and dimensionality for antigenic maps made with primary infection antisera (column 1), two dose vaccine sera (column 2), and three dose vaccine sera (column 3)**. Sera are shown as small colored squares, viruses as large circles. The grid corresponds to a two-fold dilution in the neutralization assay. Row 1 shows the antigenic map with error lines. The distance between the ends of error lines indicates the measured titer: red lines indicate that the map distance is less than measured based on the titers, blue lines when the map distance is greater than measured. Row 2 shows the difference between the table distance (estimated from the measured titer) and the fitted map distance. The dotted horizontal line shows what would be perfect a perfect fit of the data. Row 3 shows the results of dimensionality testing. Cross-validation (excluding 10% of titers as a test set in 100 independent repeats) was used to determine the optimal number of dimensions. Lower root mean squared error (RMSE) for both detectable titers (above the assay limit of detection) and undetectable (below the assay limit of detection) indicate the optimal number of dimensions for fitting the antigenic map. Row 4 shows antigenic maps made in three dimensions.

**Fig. S2. Evaluation of robustness in positioning for viruses and sera on antigenic maps made with primary infection antisera (column 1), two dose vaccine sera (column 2), and three dose vaccine sera (column 3)**. Sera are shown as open shapes, viruses as colored shapes. The grid corresponds to a two-fold dilution in the neutralization assay. Row 1 shows triangulation/coordination confidence intervals, indicating confidence in positioning of points. Each shape marks the area that the point can occupy before increasing the total map error by more than 1 antigenic unit. Row 2 shows bootstrapped maps considering titer error for the neutralization assay. The shapes correspond to the positions of points on resampled maps assuming titers have random noise added with the measured assay standard deviation of log_2_ 0.29 (1.2-fold). Row 3 shows confidence in coordination of points following bootstrapping of the sera and viruses.

**Fig. S3.** Comparison of virus positions between antigenic maps. Arrows point to virus positions from one map to another. Sera are shown as small squares, viruses as colored circles. The grid corresponds to a two-fold dilution in the neutralization assay.

**Table S1.** Characteristics of post-infection convalescent participants.

**Table S2.** Characteristics of post-vaccination uninfected participants.

**Table S3.** SARS-CoV-2 sequence data from SARS-CoV-2 infections.

**Table S4.** Commercially obtained convalescent sera.

**Table S5.** Spike mutations of commercially obtained convalescent sera.

**Table S6.** SARS-CoV2 variant spikes used in neutralization assays.

**Table S7.** Median, standard deviation, and confidence intervals for fold breadth gain for each exposure history against each measured antigen from observed data.

**Table S8.** Significance values for breadth gain comparisons

**Table S9.** Slope and Peak Location for Individual Antibody Landscapes

**Table S10.** Summary statistics of cone slope for individual landscapes grouped by exposure history

## ACKNOWLEDGMENTS

We sincerely thank the members of the EPICC COVID-19 Cohort Study Group for their many contributions in conducting the study and ensuring effective protocol operations. The authors wish to also acknowledge all who have contributed to the EPICC COVID-19 study: *Brooke Army Medical Center, Fort Sam Houston, TX: Col* J. Cowden; LTC M. Darling; S. DeLeon; Maj D. Lindholm; LTC A. Markelz; K. Mende; S. Merritt; T. Merritt; LTC N. Turner; CPT T. Wellington*. Carl R. Darnall Army Medical Center, Fort Hood, TX:* LTC S. Bazan; P.K Love*. Fort Belvoir Community Hospital, Fort Belvoir, VA:* N. Dimascio-Johnson; MAJ E. Ewers; LCDR K. Gallagher; LCDR D. Larson; A. Rutt*. Henry M. Jackson Foundation, Inc., Bethesda, MD:* P. Blair; J. Chenoweth; D. Clark*. Madigan Army Medical Center, Joint Base Lewis McChord, WA:* S. Chambers; LTC C. Colombo; R. Colombo; CAPT C. Conlon; CAPT K. Everson; COL P. Faestel; COL T. Ferguson; MAJ L. Gordon; LTC S. Grogan; CAPT S. Lis; COL C. Mount; LTC D. Musfeldt; CPT D. Odineal; LTC M. Perreault; W. Robb-McGrath; MAJ R. Sainato; C. Schofield; COL C. Skinner; M. Stein; MAJ M. Switzer; MAJ M. Timlin; MAJ S. Wood*. Naval Medical Center Portsmouth, Portsmouth, VA:* S. Banks; R. Carpenter; L. Kim; CAPT K. Kronmann; T. Lalani; LCDR T. Lee; LCDR A. Smith; R. Smith; R. Tant; T. Warkentien*. Naval Medical Center San Diego, San Diego, CA:*CDR C. Berjohn; S. Cammarata; N. Kirkland; D. Libraty; CAPT (Ret) R. Maves; CAPT (Ret) G. Utz*. Tripler Army Medical Center, Honolulu, HI:* S. Chi; LTC R. Flanagan; MAJ M. Jones; C. Lucas; LTC (Ret) C. Madar; K. Miyasato; C. Uyehara*. Uniformed Services University of the Health Sciences, Bethesda, MD:* B. Agan; L. Andronescu; A. Austin; C. Broder; CAPT T. Burgess; C. Byrne; COL (Ret.) K Chung; J. Davies; C. English; N. Epsi; C. Fox; M. Fritschlanski; M. Grother; A. Hadley; COL P. Hickey; E. Laing; LTC C. Lanteri; LTC J. Livezey; A. Malloy; R. Mohammed; C. Morales; P. Nwachukwu; C. Olsen; E. Parmelee; S. Pollett; S. Richard; J. Rozman; J. Rusiecki; E. Samuels; P. Nwachukwu; M. Tso; M. Sanchez; A. Scher; CDR M. Simons; A. Snow; K. Telu; D. Tribble; L. Ulomi*. United States Air Force School of Aerospace Medicine, Dayton, OH:* TSgt T. Chao; R. Chapleau; M. Christian; A. Fries; C. Harrington; V. Hogan; S. Huntsberger; K. Lanter; E. Macias; J. Meyer; S. Purves; K. Reynolds; J. Rodriguez; C. Starr*. United States Army Medical Research Institute of Infectious Diseases;* J Kugelman, *United States Coast Guard, Washington, DC:* CAPT J. Iskander, CDR I. Kamara*. Womack Army Medical Center, Fort Bragg, NC:* B. Barton; LTC D. Hostler; LTC J. Hostler; MAJ K. Lago; C. Maldonado; J. Mehrer*. William Beaumont Army Medical Center, El Paso, TX*: MAJ T. Hunter; J. Mejia; J. Montes; R. Mody; R. Resendez; P. Sandoval; M. Wayman*. Walter Reed National Military Medical Center, Bethesda, MD:* I. Barahona; A. Baya; A. Ganesan; MAJ N. Huprikar; B. Johnson*. Walter Reed Army Institute of Research, Silver Spring, MD*: S. Peel. The authors wish to also acknowledge the following individuals for their contributions to the PASS (IDCRP-126) COVID-19 study. *Henry M. Jackson Foundation, Inc., Bethesda, MD*: Alyssa Lindrose, Matthew Moser, Emily C. Samuels, Belinda Jackson-Thompson, Julian Davies, Luca Illinik, Mimi Sanchez, Orlando Ortega, Edward Parmelee*. NMRC-CTC (Naval Medical Research Center – Clinical Trials Center):* Santina E. Maiolatesi. Christopher A. Duplessis*. NMRC-CTC/General Dynamics*. Kathleen F. Ramsey, Anatalio E. Reyes, Yolanda Alcorta, Mimi A. Wong. We also wish to thank Drs. Douglas Pratt and Gabriel Parra (FDA) for critical review of the manuscript.

## AUTHOR CONTRIBUTIONS

a. Conceived and designed the study / experiments: WW, SL, LCK, CDW, SP

b. Acquired data / performed experiments: WW, SL, CDW, ACF, RW, EL, EG, RV, SC, MPS, DRT, THB, BKA, EM, SDP

c. Created detailed analysis plan and/or analyzed the data: WW, LCK, CDW, SL, RS, NE, ACF, FE, JOJ, SR

d. Interpreted findings: WW, LCK, CDW, SL, RS, ACF, SP, EM, EL, CB

e. Contributed resources; reagents/materials/specimens: EG, JR, DAL, KM, EE, DTL, REC, CJC, AS, TL, CMB, RCM, MUJ, RM, NH, JL, DS, MHP, GW, AG, MPS, CDW

f. Composed first draft of manuscript: WW, LCK, CDW, SL, RS, SP

g. Provided critical revisions and edits (scientific content) to provisional drafts: WW, LCK, CDW, SL, RS, NE, ACF, EM, EL, EM

h. Reviewed and approved final version for submission: All Authors

## DISCLAIMERS

The contents of this publication are the sole responsibility of the author (s) and do not necessarily reflect the views, opinions, or policies of Uniformed Services University of the Health Sciences (USUHS); the Department of Defense (DoD); the Departments of the Army, Navy, or Air Force; the Defense Health Agency, or the Henry M. Jackson Foundation for the Advancement of Military Medicine Inc. Mention of trade names, commercial products, or organizations does not imply endorsement by the U.S. Government. The investigators have adhered to the policies for protection of human subjects as prescribed in 45 CFR 46. The Emerging Leaders in Data Science Research Participation Program is administered by the Oak Ridge Institute for Science and Education through an interagency agreement between the U.S. Department of Energy (DOE) and NIAID. ORISE is managed by ORAU under DOE contract number DE-SC0014664. All opinions expressed in this paper are the author’s and do not necessarily reflect the policies and views of FDA, NIAID, DOE, or ORAU/ORISE.

