## Supplemental tables and figures for "Post-vaccination Omicron infections induce broader immunity across antigenic space than prototype mRNA COVID-19 booster vaccination or primary infection"

### Supplementary Tables

**Table S1: Characteristics of post-infection convalescent participants**

| | Overall (N=84)[a] |
|---|---|
| **Age group** | |
| <18 | 5 (6.0%) |
| 18-44 | 49 (58.3%) |
| 45-64 | 21 (25.0%) |
| 65+ | 8 (9.5%) |
| Missing | 1 (1.2%) |
| **Gender** | |
| Female | 33 (39.3%) |
| Male | 51 (60.7%) |
| **Race/ Ethnicity** | |
| Black | 10 (11.9%) |
| Hispanic or Latino | 20 (23.8%) |
| Others | 8 (9.5%) |
| White | 45 (53.6%) |
| Missing | 1 (1.2%) |
| **Severity of initial infection** | |
| Hospitalized | 26 (31.0%) |
| Outpatient | 56 (66.7%) |
| Missing | 2 (2.4%) |
| **Charlson comorbidity index** | |
| 0 | 46 (56.8%) |
| 1-2 | 21 (25.9%) |
| 3-4 | 9 (11.1%) |
| >5 | 5 (6.2%) |
| Missing | 3 (3.6%) |
| **Days between infection symptom onset and sera sample collection** | |
| Mean ± SD (range) | 23.9 ± 10.7 (3.0 - 51.0) |
| **Infecting genotype[b]** | |
| BA.1.1 | 12 (14.3%) |
| BA.1 | 11 (13.1%) |
| B.1 | 10 (11.9%) |
| B.1.2 | 8 (9.5%) |
| B.1.617.2 | 4 (4.8%) |
| B.1.1.207 | 3 (3.6%) |
| B.1.1.7 | 3 (3.6%) |
| AY.100 | 2 (2.4%) |

| | |
|---|---|
| AY.14 | 2 (2.4%) |
| AY.25 | 2 (2.4%) |
| B.1.429 | 2 (2.4%) |
| AY.119 | 1 (1.2%) |
| AY.25.1 | 1 (1.2%) |
| AY.3 | 1 (1.2%) |
| AY.44 | 1 (1.2%) |
| AY.47 | 1 (1.2%) |
| AY.62 | 1 (1.2%) |
| AY.74 | 1 (1.2%) |
| B.1.1.318 | 1 (1.2%) |
| B.1.1.519 | 1 (1.2%) |
| B.1.243 | 1 (1.2%) |
| B.1.351 | 1 (1.2%) |
| B.1.526 | 1 (1.2%) |
| P.1.10 | 1 (1.2%) |
| Q.4 | 1 (1.2%) |
| Omicron[c] | 1 (1.2%) |
| Presumptive Beta[d] | 1 (1.2%) |
| Presumptive Delta[e] | 2 (2.4%) |
| Presumptive Pre-Delta[e] | 7 (8.3%) |
| **Study Group** | |
| EPICC | 81 (96.4%) |
| PASS | 2 (2.4%) |
| CBER | 1 (1.2%) |
| **Primary vaccine** | |
| mRNA-1273 (Moderna) | 5 (6.0%) |
| BNT162b2 (Pfizer) | 32 (38.1%) |
| Unvaccinated | 47 (56.0%) |
| **Vaccination status by time of infection and sera sampling** | |
| Unvaccinated | 47 (56.0%) |
| Vaccinated, not boosted | 27 (32.1%) |
| Boosted | 10 (11.9%) |

[a]*Sample collected from EPICC, PASS and CBER*

[b]*Genotypes assigned based on Pango 4.0.6*

[c]*Genotype assigned based on Nextclade*

[d]*Sera from a traveler who had moderate-severe COVID-19 (outpatient) in the Republic of South Africa during the peak of the Beta (B.1.351) wave in January 2021.*

[e]*Inferred by date of collection*

**Table S2: Characteristics of post-vaccination uninfected participants**

| | Overall (N=39)[a] |
|---|---|
| **Age** | |
| 18-44 | 19 (48.5%) |
| 45-64 | 19 (48.5%) |
| 65+ | 1 (3.0%) |
| **Gender** | |
| Female | 25 (64.0%) |
| Male | 14 (36.0%) |
| **Race** | |
| White | 26 (67.0%) |
| Asian | 8 (21.0%) |
| Black | 4 (10.0%) |
| Multiracial | 1 (3.0%) |
| **Occupation** | |
| Nurse | 11 (28.0%) |
| Physician | 11 (28.0%) |
| Physical/Occupational/Recreational Therapist | 9 (23.0%) |
| Lab personnel | 3 (8.0%) |
| Medical Technician/Support Assistant | 3 (8.0%) |
| Social Worker | 1 (3.0%) |
| Psychologist | 1 (3.0%) |
| **Time between 2nd vaccine and sera sample collection (days)** | |
| Mean ± SD (range) | 30.5 ± 3.1 (22 - 42) |
| **Time between booster dose and sera sample collection (days)** | |
| Mean ± SD (range) | 42.6 ± 16.9 (7 - 93) |

[a]*Samples collected from the PASS study*

**Table S3: SARS-CoV-2 sequence data from SARS-CoV-2 infections**

| Sample# | Pangolin 4.0.6 | Spike Substitutions | Spike Deletions | Accession |
|---|---|---|---|---|
| Conv-01 | BA.1.1 | S:A67V,S:D80N,S:T95I,S:Y145D,S:L212I,S:G339D,S:R346K,S:S371L,S:S373P,S:S375F,S:K417N,S:N440K,S:G446S,S:S477N,S:T478K,S:E484A,S:Q493R,S:G496S,S:Q498R,S:N501Y,S:Y505H,S:T547K,S:D614G,S:H655Y,S:N679K,S:P681H,S:N764K,S:D796Y,S:N856K,S:Q954H,S:N969K,S:L981F | S:H69-,S:V70-,S:G142-,S:V143-,S:Y144-,S:N211- | ON897697 |
| Conv-02 | BA.1.1 | S:A67V,S:T95I,S:Y145D,S:L212I,S:F318S,S:G339D,S:R346K,S:S371L,S:S373P,S:S375F,S:K417N,S:N440K,S:G446S,S:S477N,S:T478K,S:E484A,S:Q493R,S:G496S,S:Q498R,S:N501Y,S:Y505H,S:T547K,S:D614G,S:H655Y,S:N679K,S:P681H,S:N764K,S:D796Y,S:N856K,S:Q954H,S:N969K,S:L981F | S:H69-,S:V70-,S:G142-,S:V143-,S:Y144-,S:N211- | ON897698 |
| Conv-03 | BA.1 | S:A67V,S:T95I,S:Y145D,S:L212I,S:G339D,S:N679K,S:P681H,S:N764K,S:D796Y,S:N856K,S:Q954H,S:N969K,S:L981F | S:H69-,S:V70-,S:G142-,S:V143-,S:Y144-,S:N211- | ON897699 |

| Sample# | Pangolin 4.0.6 | Spike Substitutions | Spike Deletions | Accession |
|---|---|---|---|---|
| Conv-04 | BA.1.1.18 | S:A67V,S:T95I,S:Y145D,S:L212I,S:G339D,S:R346K,S:S371L,S:S373P,S:S375F,S:K417N,S:N440K,S:G446S,S:S477N,S:T478K,S:E484A,S:Q493R,S:G496S,S:Q498R,S:N501Y,S:Y505H,S:T547K,S:D614G,S:H655Y,S:N679K,S:P681H,S:N764K,S:D796Y,S:N856K,S:Q954H,S:N969K,S:L981F | S:H69-,S:V70-,S:G142-,S:V143-,S:Y144-,S:N211- | ON897700 |
| Conv-05 | BA.1.1 | S:A67V,S:T95I,S:Y145D,S:L212I,S:G339D,S:R346K,S:S371L,S:S373P,S:S375F,S:K417N,S:N440K,S:G446S,S:S477N,S:T478K,S:E484A,S:Q493R,S:G496S,S:Q498R,S:N501Y,S:Y505H,S:T547K,S:D614G,S:H655Y,S:N679K,S:P681H,S:N764K,S:D796Y,S:N856K,S:Q954H,S:N969K,S:L981F | S:H69-,S:V70-,S:G142-,S:V143-,S:Y144-,S:N211- | ON897701 |
| Conv-06 | BA.1.1 | S:A67V,S:T95I,S:Y145D,S:L212I,S:G339D,S:R346K,S:S371L,S:S373P,S:S375F,S:K417N,S:N440K,S:G446S,S:S477N,S:T478K,S:E484A,S:Q493R,S:G496S,S:Q498R,S:N501Y,S:Y505H,S:T547K,S:D614G,S:H655Y,S:N679K,S:P681H,S:N764K,S:D796Y,S:N856K,S:Q954H,S:N969K,S:L981F | S:H69-,S:V70-,S:G142-,S:V143-,S:Y144-,S:N211- | ON897704 |

| Sample# | Pangolin 4.0.6 | Spike Substitutions | Spike Deletions | Accession |
|---|---|---|---|---|
| Conv-07 | BA.1.1 | S:A67V,S:T95I,S:Y145D,S:L212I,S:G339D,S:R346K,S:S371L,S:S373P,S:S375F,S:K417N,S:N440K,S:G446S,S:S477N,S:T478K,S:E484A,S:Q493R,S:G496S,S:Q498R,S:N501Y,S:Y505H,S:T547K,S:D614G,S:H655Y,S:N679K,S:P681H,S:N764K,S:D796Y,S:N856K,S:Q954H,S:N969K,S:L981F | S:H69-,S:V70-,S:G142-,S:V143-,S:Y144-,S:N211- | ON897705 |
| Conv-08 | BA.1.1 | S:A67V,S:T95I,S:Y145D,S:L212I,S:G339D,S:R346K,S:S371L,S:S373P,S:S375F,S:K417N,S:N440K,S:G446S,S:S477N,S:T478K,S:E484A,S:Q493R,S:G496S,S:Q498R,S:N501Y,S:Y505H,S:T547K,S:D614G,S:H655Y,S:N679K,S:P681H,S:N764K,S:D796Y,S:N856K,S:Q954H,S:N969K,S:L981F | S:H69-,S:V70-,S:G142-,S:V143-,S:Y144-,S:N211- | ON897706 |
| Conv-09 | BA.1.1 | S:A67V,S:T95I,S:Y145D,S:L212I,S:G339D,S:R346K,S:S371L,S:S373P,S:S375F,S:K417N,S:N440K,S:G446S,S:S477N,S:T478K,S:E484A,S:Q493R,S:G496S,S:Q498R,S:N501Y,S:Y505H,S:T547K,S:D614G,S:H655Y,S:N679K,S:P681H,S:N764K,S:D796Y,S:N856K,S:Q954H,S:N969K,S:L981F | S:H69-,S:V70-,S:G142-,S:V143-,S:Y144-,S:N211- | ON897707 |

| Sample# | Pangolin 4.0.6 | Spike Substitutions | Spike Deletions | Accession |
|---|---|---|---|---|
| Conv-10 | BA.1.1 | S:A67V,S:T95I,S:Y145D,S:L212I,S:G339D,S:R346K,S:S371L,S:S373P,S:S375F,S:K417N,S:N440K,S:G446S,S:S477N,S:T478K,S:E484A,S:Q493R,S:G496S,S:Q498R,S:N501Y,S:Y505H,S:T547K,S:D614G,S:H655Y,S:N679K,S:P681H,S:N764K,S:D796Y,S:N856K,S:Q954H,S:N969K,S:L981F | S:H69-,S:V70-,S:G142-,S:V143-,S:Y144-,S:N211- | ON897708 |
| Conv-11 | BA.1.1 | S:A67V,S:T95I,S:Y145D,S:L212I,S:G339D,S:R346K,S:S371L,S:S373P,S:S375F,S:K417N,S:N440K,S:G446S,S:S477N,S:T478K,S:E484A,S:Q493R,S:G496S,S:Q498R,S:N501Y,S:Y505H,S:T547K,S:D614G,S:H655Y,S:N679K,S:P681H,S:N764K,S:D796Y,S:N856K,S:Q954H,S:N969K,S:L981F | S:H69-,S:V70-,S:G142-,S:V143-,S:Y144-,S:N211- | ON897709 |
| Conv-12 | BA.1.1 | S:A67V,S:T95I,S:Y145D,S:L212I,S:G339D,S:R346K,S:S371L,S:S373P,S:S375F,S:K417N,S:N440K,S:G446S,S:S477N,S:T478K,S:E484A,S:Q493R,S:G496S,S:Q498R,S:N501Y,S:Y505H,S:T547K,S:D614G,S:H655Y,S:N679K,S:P681H,S:N764K,S:D796Y,S:N856K,S:Q954H,S:N969K,S:L981F | S:H69-,S:V70-,S:G142-,S:V143-,S:Y144-,S:N211- | ON897710 |

| Sample# | Pangolin 4.0.6 | Spike Substitutions | Spike Deletions | Accession |
|---|---|---|---|---|
| Conv-13 | BA.1.1 | S:A67V,S:T95I,S:Y145D,S:L212I,S:G339D,S:R346K,S:S371L,S:S373P,S:S375F,S:K417N,S:N440K,S:G446S,S:S477N,S:T478K,S:E484A,S:Q493R,S:G496S,S:Q498R,S:N501Y,S:Y505H,S:T547K,S:D614G,S:H655Y,S:N679K,S:P681H,S:N764K,S:D796Y,S:N856K,S:Q954H,S:N969K,S:L981F | S:H69-,S:V70-,S:G142-,S:V143-,S:Y144-,S:N211- | SAMN29442565 |
| Conv-14 | BA.1.21 | S:A67V,S:T95I,S:Y145D,S:L212I,S:G339D,S:S371L,S:S373P,S:S375F,S:K417N,S:N440K,S:G446S,S:S477N,S:T478K,S:E484A,S:Q493R,S:G496S,S:Q498R,S:N501Y,S:Y505H,S:T547K,S:D614G,S:E654K,S:H655Y,S:N679K,S:P681H | S:H69-,S:V70-,S:G142-,S:V143-,S:Y144-,S:N211- | ON897711 |
| Conv-15 | BA.1 | S:A67V,S:T95I,S:Y145D,S:L212I,S:G339D,S:S371L,S:S373P,S:S375F,S:K417N,S:N440K,S:G446S,S:S477N,S:T478K,S:E484A,S:Q493R,S:G496S,S:Q498R,S:N501Y,S:Y505H,S:T547K,S:D614G,S:H655Y,S:N679K,S:P681H | S:H69-,S:V70-,S:G142-,S:V143-,S:Y144-,S:N211- | SAMN29442568 |

| Sample# | Pangolin 4.0.6 | Spike Substitutions | Spike Deletions | Accession |
|---|---|---|---|---|
| Conv-16 | BA.1.18 | S:A67V,S:T95I,S:Y145D,S:L212I,S:G339D,S:S371L,S:S373P,S:S375F,S:K417N,S:N440K,S:G446S,S:S477N,S:T478K,S:E484A,S:Q493R,S:G496S,S:Q498R,S:N501Y,S:Y505H,S:T547K,S:D614G,S:H655Y,S:N679K,S:P681H,S:N764K,S:D796Y,S:N856K,S:Q954H,S:N969K,S:L981F | S:H69-,S:V70-,S:G142-,S:V143-,S:Y144-,S:N211- | ON897713 |
| Conv-17 | BA.1.18 | S:A67V,S:T95I,S:Y145D,S:L212I,S:G339D,S:S371L,S:S373P,S:S375F,S:K417N,S:N440K,S:G446S,S:S477N,S:T478K,S:E484A,S:Q493R,S:G496S,S:Q498R,S:N501Y,S:Y505H,S:T547K,S:D614G,S:H655Y,S:N679K,S:P681H,S:N764K,S:D796Y,S:N856K,S:Q954H,S:N969K,S:L981F | S:H69-,S:V70-,S:G142-,S:V143-,S:Y144-,S:N211- | ON897714 |
| Conv-18 | BA.1.15 | S:A67V,S:T95I,S:Y145D,S:L212I,S:G339D,S:S371L,S:S373P,S:S375F,S:K417N,S:N440K,S:G446S,S:S477N,S:T478K,S:E484A,S:Q493R,S:G496S,S:Q498R,S:N501Y,S:Y505H,S:T547K,S:D614G,S:H655Y,S:N679K,S:P681H,S:N764K,S:D796Y,S:N856K,S:Q954H,S:N969K,S:L981F | S:H69-,S:V70-,S:G142-,S:V143-,S:Y144-,S:N211- | ON897715 |

| Sample# | Pangolin 4.0.6 | Spike Substitutions | Spike Deletions | Accession |
|---|---|---|---|---|
| Conv-19 | BA.1.15 | S:A67V,S:T95I,S:Y145D,S:L212I,S:G339D,S:S371L,S:S373P,S:S375F,S:K417N,S:N440K,S:G446S,S:S477N,S:T478K,S:E484A,S:Q493R,S:G496S,S:Q498R,S:N501Y,S:Y505H,S:T547K,S:D614G,S:H655Y,S:N679K,S:P681H,S:N764K,S:D796Y,S:N856K,S:Q954H,S:N969K,S:L981F | S:H69-,S:V70-,S:G142-,S:V143-,S:Y144-,S:N211- | ON897718 |
| Conv-20 | BA.1.18 | S:A67V,S:T95I,S:Y145D,S:L212I,S:G339D,S:S371L,S:S373P,S:S375F,S:K417N,S:N440K,S:G446S,S:S477N,S:T478K,S:E484A,S:Q493R,S:G496S,S:Q498R,S:N501Y,S:Y505H,S:T547K,S:D614G,S:H655Y,S:N679K,S:P681H,S:N764K,S:D796Y,S:N856K,S:Q954H,S:N969K,S:L981F | S:H69-,S:V70-,S:G142-,S:V143-,S:Y144-,S:N211- | ON897720 |
| Conv-21 | BA.1.15.2 | S:A67V,S:T95I,S:Y145D,S:L212I,S:V320I,S:G339D,S:S371L,S:S373P,S:S375F,S:K417N,S:N440K,S:G446S,S:S477N,S:T478K,S:E484A,S:Q493R,S:G496S,S:Q498R,S:N501Y,S:Y505H,S:T547K,S:D614G,S:Q628K,S:H655Y,S:N679K,S:P681H,S:N764K,S:D796Y,S:N856K,S:Q954H,S:N969K,S:L981F | S:H69-,S:V70-,S:G142-,S:V143-,S:Y144-,S:N211- | ON897722 |

| Sample# | Pangolin 4.0.6 | Spike Substitutions | Spike Deletions | Accession |
|---|---|---|---|---|
| Conv-22 | BA.1.1 | S:A67V,S:T95I,S:Y145D,S:P209S,S:L212I,S:G339D,S:R346K,S:S371L,S:S373P,S:S375F,S:K417N,S:N440K,S:G446S,S:S477N,S:T478K,S:E484A,S:Q493R,S:G496S,S:Q498R,S:N501Y,S:Y505H,S:T547K,S:D614G,S:H655Y,S:N679K,S:P681H | S:H69-,S:V70-,S:G142-,S:V143-,S:Y144-,S:N211- | ON897725 |
| Conv-23 | B.1 | S:D614G | | OM000280 |
| Conv-24 | B.1 | S:D614G | | OM000281 |
| Conv-25 | B.1 | S:D614G | | OM000284 |
| Conv-26 | B.1 | S:D614G | | OM000285 |
| Conv-27 | B.1 | S:D614G | | OM000282 |
| Conv-28 | B.1.2 | S:D614G | | OM000296 |
| Conv-29 | B.1 | S:D614G | | OM000286 |
| Conv-30 | B.1 | S:D614G | | OM000279 |
| Conv-31 | B.1 | S:D614G | | OM000288 |
| Conv-32 | B.1 | S:D614G | | OM000287 |
| Conv-33 | B.1 | S:D614G | | OM000283 |
| Conv-34 | B.1.2 | S:D614G | | OM000297 |
| Conv-35 | B.1.2 | S:D614G | | OM000299 |
| Conv-36 | B.1.2 | S:D614G | | OM000298 |
| Conv-37 | B.1.2 | S:D614G | | OM000294 |
| Conv-38 | B.1.1.207 | S:D614G,S:P681H | | ON897726 |
| Conv-39 | B.1.1.207 | S:E484K,S:D614G,S:P681H | | ON897727 |
| Conv-40 | B.1.1.207 | S:E484K,S:D614G,S:P681H | | ON897728 |
| Conv-41 | B.1.2 | S:G257D,S:D614G | | OM000295 |

| Sample# | Pangolin 4.0.6 | Spike Substitutions | Spike Deletions | Accession |
|---|---|---|---|---|
| Conv-42 | P.1.10 | S:L18F,S:T20N,S:P26S,S:D138Y,S:R190S,S:K417T,S:E484K,S:N501Y,S:D614G,S:H655Y,S:S704L,S:T1027I,S:A1078S,S:V1176F | | ON897729 |
| Conv-43 | B.1.1.519 | S:L5F,S:T478K,S:D614G,S:P681H,S:T732A | | ON897730 |
| Conv-44 | B.1.526 | S:L5F,S:T95I,S:D253G,S:E484K,S:D614G,S:A701V | | ON897731 |
| Conv-45 | B.1.1.7 | S:N501Y,S:A570D,S:D614G,S:P681H,S:T716I,S:Q836*,S:S982A,S:D1118H | S:H69-,S:V70-,S:Y144 | OM000289 |
| Conv-46 | B.1.1.7 | S:N501Y,S:A570D,S:D614G,S:P681H,S:T716I,S:S982A,S:D1118H | S:H69-,S:V70-,S:Y144- | OM000292 |
| Conv-47 | B.1.1.7 | S:N501Y,S:A570D,S:D614G,S:P681H,S:T716I,S:S982A,S:D1118H,S:K1191N | S:H69-,S:V70-,S:Y144- | OM000291 |
| Conv-48 | B.1.429 | S:S13I,S:T95I,S:W152C,S:L452R,S:D614G | | ON897732 |
| Conv-49 | B.1.429 | S:S13I,S:W152C,S:L452R,S:D614G | | ON897733 |
| Conv-50 | B.1.617.2 | S:T19R,S:G142D,S:M153I,S:R158G,S:A222V,S:L452R,S:T478K,S:D614G,S:P681R,S:D950N | S:E156-,S:F157- | OM000277 |
| Conv-51 | AY.74 | S:T19R,S:G142D,S:R158G,S:A222V,S:L452R,S:T478K,S:D614G,S:P681R,S:D950N | S:E156-,S:F157- | OM000262 |

| Sample# | Pangolin 4.0.6 | Spike Substitutions | Spike Deletions | Accession |
|---|---|---|---|---|
| Conv-52 | AY.62 | S:T19R,S:G142D,S:R158G,S:A222V,S:L452R,S:T478K,S:D614G,S:P681R,S:G946V,S:D950N | S:E156-,S:F157- | OM000270 |
| Conv-53 | AY.47 | S:T19R,S:G142D,S:R158G,S:A222V,S:V289I,S:L452R,S:T478K,S:D614G,S:P681R,S:D950N | S:E156-,S:F157- | OM000264 |
| Conv-54 | AY.25.1 | S:T19R,S:G142D,S:R158G,S:L452R,S:T478K,S:D614G,S:P681R,S:D950N | S:E156-,S:F157- | OM000271 |
| Conv-55 | AY.14 | S:T19R,S:G142D,S:R158G,S:L452R,S:T478K,S:D614G,S:P681R,S:D950N | S:E156-,S:F157- | OM000267 |
| Conv-56 | AY.14 | S:T19R,S:G142D,S:R158G,S:L452R,S:T478K,S:D614G,S:P681R,S:D950N | S:E156-,S:F157- | OM000266 |
| Conv-57 | AY.3 | S:T19R,S:G142D,S:R158G,S:L452R,S:T478K,S:D614G,S:P681R,S:D950N | S:E156-,S:F157- | OM000274 |
| Conv-58 | B.1.617.2 | S:T19R,S:G142D,S:R158G,S:L452R,S:T478K,S:D614G,S:P681R,S:D950N,S:V1264L | S:E156-,S:F157- | OM000275 |
| Conv-59 | B.1.617.2 | S:T19R,S:K77T,S:G142D,S:R158G,S:G181V,S:L452R,S:T478K,S:D614G,S:A653V,S:P681R,S:D950N | S:E156-,S:F157- | OM000265 |

| Sample# | Pangolin 4.0.6 | Spike Substitutions | Spike Deletions | Accession |
|---|---|---|---|---|
| Conv-60 | B.1.617.2 | S:T19R,S:K77T,S:G142D,S:R158G,S:G181V,S:L452R,S:T478K,S:D614G,S:A653V,S:P681R,S:D950N | S:E156-,S:F157- | OM311576 |
| Conv-61 | AY.25 | S:T19R,S:S112L,S:G142D,S:R158G,S:L452R,S:T478K,S:D614G,S:P681R,S:D950N | S:E156-,S:F157- | OM000263 |
| Conv-62 | AY.25 | S:T19R,S:S112L,S:G142D,S:R158G,S:L452R,S:T478K,S:D614G,S:P681R,S:D950N | S:E156-,S:F157- | OM000268 |
| Conv-63 | AY.44 | S:T19R,S:T22I,S:G142D,S:R158G,S:L452R,S:T478K,S:D614G,S:P681R,S:D950N | S:E156-,S:F157- | OM000272 |
| Conv-64 | AY.100 | S:T19R,S:T95I,S:G142D,S:R158G,S:L452R,S:T478K,S:D614G,S:P681R,S:D950N | S:E156-,S:F157- | OM000276 |
| Conv-65 | AY.119 | S:T19R,S:T95I,S:G142D,S:R158G,S:L452R,S:T478K,S:D614G,S:P681R,S:D950N | S:E156-,S:F157- | OM000273 |
| Conv-66 | AY.100 | S:T19R,S:T95I,S:G142D,S:R158G,S:L452R,S:T478K,S:D614G,S:P681R,S:D950N,S:G1124V | S:E156-,S:F157- | OM000269 |
| Conv-67 | Q.4 | S:N501Y,S:A570D,S:D614G,S:P681R,S:T716I,S:S982A,S:D1118H | S:H69-,S:V70-,S:Y144- | ON897734 |
| Conv-68 | B.1.1.318 | S:D614G | | ON897735 |
| Conv-69 | B.1.2 | S:D614G | | ON897736 |
| Conv-70 | B.1.2 | S:D614G | | ON897737 |

| Sample# | Pangolin 4.0.6 | Spike Substitutions | Spike Deletions | Accession |
|---|---|---|---|---|
| Conv-71 | B.1.243 | S:L5F,S:D614G,S:P681H,S:T859I | | ON897738 |
| Conv-72 | BA.1 | | | ON897739 |
| Conv-73 | Omicron[a] | | | |
| Conv-74 | B.1.351[b] | | | |
| Conv-75 | Presumptive Pre-Delta[c] | | | |
| Conv-76 | Presumptive Pre-Delta[c] | | | |
| Conv-77 | Presumptive Pre-Delta[c] | | | |
| Conv-78 | Presumptive Pre-Delta[c] | | | |
| Conv-79 | Presumptive Pre-Delta[c] | | | |
| Conv-80 | Presumptive Pre-Delta[c] | | | |
| Conv-81 | Presumptive Pre-Delta[c] | | | |

[a]*Genotype determined by Nextclade*

[b]*The SARS-CoV-2 sequence from this individual was not available*

[c]*Genotype and sequence were not available*

**Table S4: Commercially obtained convalescent sera**

| | Overall (N=31) |
|---|---|
| **Gender** | |
| Female | 10 (32.3%) |
| Male | 21 (67.7%) |
| **Infecting genotype** | |
| B.1 | 2 (6.5%) |
| B.1.1.7 | 7 (22.6%) |
| B.1.2 | 2 (6.5%) |
| B.1.234 | 1 (3.2%) |
| B.1.429 | 3 (9.7%) |
| B.1.577 | 1 (3.2%) |
| C.11 | 1 (3.2%) |
| C.37 | 10 (32.3%) |
| P.1 | 4 (12.9%) |
| **Time between infection symptom onset and sera sample collection** | |
| Mean ± SD (range) | 9.6 ± 12.6 (2.0 - 59.0) |
| **Commercial Sera** | |
| Boca-1 | 10 (32.3%) |
| Boca-2 | 10 (32.3%) |
| Boca-3 | 11 (35.5%) |

**Table S5: Spike mutations of Commercially obtained convalescent sera**

| Specimen ID | Infecting Genotype | Spike mutations |
|---|---|---|
| 738741 | B.1 | S:S13I;S:Q52R;S:A67V;S:L452R |
| 743259 | B.1 | D614G |
| 718055 | B.1.2 | D614G |
| 718057 | B.1.2 | S24L;S:D614G |
| 743136 | B.1.234 | D614G |
| 743256 | B.1.429 | S:W152C;S:L452R;S:D614G |
| 743257 | B.1.429 | S:L452R;S:D614G |
| 743264 | B.1.429 | S:S13I;S:W152C;S:L452R;S:D614G |
| 719166 | B.1.577 | D614G |
| 743255 | C.11 | S:L452R;S:D614G |
| D000113656 | B.1.1.7 | S:N501Y;S:A570D;S:D614G;S:P681H;S:T716I;S:S982A;S:D1118H |
| D000113657 | B.1.1.7 | S:N501Y;S:A570D;S:D614G;S:P681H;S:T716I;S:S982A;S:D1118H |
| D000113667 | B.1.1.7 | S:V433F;S:A570D |
| D000113669 | B.1.1.7 | S:N501Y;S:A570D;S:D614G;S:P681H;S:T716I;S:S982A;S:D1118H |
| D000113675 | B.1.1.7 | S:V193L;S:W436*;S:V510L;S:A1020S;S:P1079S;S:D1118H |
| D000113694 | B.1.1.7 | S:N501Y;S:A570D;S:D614G;S:P681H;S:T716I;S:S982A;S:D1118H;S:P1263L |
| D000117099 | C.37 | S:G75V;S:T76I;S:R246N;S:L452Q;S:F490S;S:D614G;S:T859N |
| D000117104 | C.37 | S:G75V;S:T76I;S:R246N;S:L452Q;S:F490S;S:D614G;S:T859N |
| D000117136 | C.37 | S:G75V;S:T76I;S:R246N;S:D442Y;S:L452Q;S:F490S;S:Q580X;S:T581X;S:D614G;S:I714V |

| Specimen ID | Infecting Genotype | Spike mutations |
|---|---|---|
| D000117154 | C.37 | S:G75V;S:T76I;S:R246N;S:L452Q;S:F490S;S:D614G;S:T859N |
| D00011366 | B.1.1.7 | S:N501Y;S:A570D;S:P681H;S:T716I;S:S982A;S:D1118H |
| D00012307 | P.1 | S:E484K;S:N501Y;S:D614G;S:H655Y;S:Q677R;S:T1027I;S:V1176F |
| D00012308 | C.37 | S:G75V;S:T76I;S:R246N; S:S247-;S:Y248-;S:L249-;S:T250-;S:P251-;S:G252-;S:D253-;S:L452Q;S:D614G;S:Q675H;S:I720V;S:T859N;S:P863L;S:Q1180 |
| D00012308 | C.37 | S:G75V;S:T76I;S:R246N; S:S247-;S:Y248-;S:L249-;S:T250-;S:P251-;S:G252-;S:D253-;S:L452Q;S:F490S;S:D614G;S:I714V;S:T859N;S:G1219C |
| D00012308 | C.37 | S:W64-;S:H66-;S:A67-;S:I68-; S:T63X;S:F65X;S:H69-;S:V70-;S:S71-;S:G72-;S:T73-;S:N74-;S:G75-;S:T76-; S:D138Y;S:R246N;S:S247-;S:Y248-;S:L249-;S:T250-;S:P251-;S:G252-;S:D253- S:L452Q;S:F490S;S:D614G;S:T859N |
| D000123091 | C.37 | S:G75V;S:T76I; S:S247-;S:Y248-;S:L249-;S:T250-;S:P251-;S:G252-;S:D253- S:R246N;S:L452Q;S:D614G;S:T859N |
| D000123095 | C.37 | S:G75V;S:T76I; S:S247-;S:Y248-;S:L249-;S:T250-;S:P251-;S:G252-;S:D253-;S:R246N;S:L452Q;S:F490S;S:D614G;S:T859N |
| D000123099 | C.37 | ;S:G75V;S:T76I; S:S247-;S:Y248-;S:L249-;S:T250-;S:P251-;S:G252-;S:D253-;S:R246N;S:L452Q;S:F490S;S:D614G;S:T859N |
| D000123108 | P.1 | S:D138Y;S:R190S;S:K417T;S:E484K;S:N501Y;S:D614G;S:H655Y;S:Q677R;S:T1027I;S:V1176F |
| D000123182 | P.1 | S:L18F;S:T20N;S:P26S;S:D138Y;S:R190S;S:K417T;S:E484K;S:N501Y;S:D614G;S:H655Y;S:T1027I;S:V1176F |
| D000123194 | P.1 | S:D138Y;S:K417T;S:E484K;S:N501Y;S:D614G;S:H655Y;S:T1027I;S:V1176F |

**Table S6: SARS-CoV2 variant spikes used in neutralization**

| SARS-CoV-2 Variants | Mutations in spike (compared to wild type Wuhan-Hu-1) |
|---|---|
| WT_D614G | D614G |
| N501Y_D614G | N501Y, D614G |
| E484K_D614G | E484K, D614G |
| T478K_D614G | T478K, D614G |
| L452R_D614G | L452R, D614G |
| R346K_D614G | R346K, D614G |
| K417N_D614G | K417N, D614G |
| N501Y_E484K_D614G | N501Y, E484K, D614G |
| R.1 | W152L, E484K, D614G, G769V |
| Delta (B.1.617.2) | T19R, G142D, E156-, F157-, R158G, L452R, T478K, D614G, P681R, D950N |
| Alpha (B.1.1.7) | H69−, V70−, Y144−, N501Y, A570D, D614G, P681H, T716I, S982A, D1118H |
| Beta (B.1.351) | D80A, D215G, L241−, L242−, A243−, K417N, E484K, N501Y, D614G, A701V |
| Epsilon (B.1.427) | S13I, W152C, L452R, D614G |
| Iota (B.1.526) | L5F, T95I, D253G, E484K, D614G, A701V |
| Mu (B.1.621) | T95I, Y144T, Y144S, ins145N, R346K, E484K, N501Y, D614G, P681H, D950N |
| Lambda (C.37) | G75V, T76I, R246-, S247-, Y248-, L249-, T250-, P251-, G252-, D253N, L452Q, F490S, D614G, T859N |
| Gamma (P.1) | L18F, T20N, P26S, D138Y, R190S, K417T, E484K, N501Y, D614G, H655Y, T1027I, V1176F |
| Omicron (BA.1) | A67V, del69-70, T95I, del142-144, Y145D, del211, L212I, ins214EPE, G339D, S371L, S373P, S375F, K417N, N440K, G446S, S477N, T478K, E484A, Q493R, G496S, Q498R, N501Y, Y505H, T547K, D614G, H655Y, N679K, P681H, N764K, D796Y, N856K, Q954H, N969K, L981F |
| Omicron (BA.1.1) | A67V, del69-70, T95I, del142-144, Y145D, del211, L212I, ins214EPE, G339D, R346K, S371L, S373P, S375F, K417N, N440K, G446S, S477N, T478K, E484A, Q493R, G496S, Q498R, N501Y, Y505H, T547K, D614G, H655Y, N679K, P681H, N764K, D796Y, N856K, Q954H, N969K, L981F |
| Omicron (BA.2) | T19I, del24-26, A27S, G142D, V213G, G339D, S371F, S373P, S375F, S376A, D405N, R408S, K417N, N440K, S477N, T478K, E484A, |

| | |
|---|---|
| | Q493R, Q498R, N501Y, Y505H, D614G, H655Y, N679K, P681H, N764K, D796Y, Q954H, N969K |
| Omicron (BA.2.12.1) | T19I, del24-26, A27S, G142D, V213G, G339D, S371F, S373P, S375F, S376A, D405N, R408S, K417N, N440K, L452Q, S477N, T478K, E484A, Q493R, Q498R, N501Y, Y505H, D614G, H655Y, N679K, P681H, S704L, N764K, D796Y, Q954H, N969K |
| Omicron (BA.4/BA.5) | T19I, del24-26, A27S, del69-70, G142D, V213G, G339D, S371F, S373P, S375F, T376A, D405N, R408S, K417N, N440K, L452R, S477N, T478K, E484A, F486V, Q498R, N501Y, Y505H, D614G, H655Y, N679K, P681H, N764K, D796Y, Q954H, N969K |

**Table S7: Median, standard deviation, and confidence intervals for fold breadth gain for each exposure history against each measured antigen from observed data**

| Measured Antigen | Exposure History | No. of Sera | Median Fold Breadth Gain | Std Dev | 2.5% Quantile | 97.5% Quantile | Standard Error |
|---|---|---|---|---|---|---|---|
| WT_D614G | 2-Dose Omicron BA.1/BA.1.1 PVI | 11 | 1.00 | 0.02 | 0.94 | 1.00 | 0.01 |
| WT_D614G | 3-Dose Omicron BA.1/BA.1.1 PVI | 10 | 1.00 | 0.00 | 1.00 | 1.00 | 0.00 |
| WT_D614G | 3 Dose Vaccine Only | 39 | 1.00 | 0.00 | 1.00 | 1.00 | 0.00 |
| Beta_B.1.351 | 2-Dose Omicron BA.1/BA.1.1 PVI | 11 | 5.96 | 2.16 | 2.92 | 10.08 | 0.65 |
| Beta_B.1.351 | 3-Dose Omicron BA.1/BA.1.1 PVI | 10 | 5.45 | 1.98 | 3.38 | 9.59 | 0.63 |
| Beta_B.1.351 | 3 Dose Vaccine Only | 39 | 5.50 | 2.85 | 1.58 | 12.06 | 0.46 |
| Delta_B.1.617.2 | 2-Dose Omicron BA.1/BA.1.1 PVI | 11 | 1.54 | 0.63 | 1.22 | 2.92 | 0.19 |
| Delta_B.1.617.2 | 3-Dose Omicron BA.1/BA.1.1 PVI | 10 | 1.75 | 0.28 | 1.48 | 2.28 | 0.09 |
| Delta_B.1.617.2 | 3 Dose Vaccine Only | 39 | 0.98 | 0.22 | 0.65 | 1.50 | 0.04 |
| Mu_B.1.621 | 2-Dose Omicron BA.1/BA.1.1 PVI | 11 | 4.35 | 1.38 | 2.86 | 6.95 | 0.41 |
| Mu_B.1.621 | 3-Dose Omicron BA.1/BA.1.1 PVI | 10 | 4.10 | 1.70 | 2.68 | 7.82 | 0.54 |
| Mu_B.1.621 | 3 Dose Vaccine Only | 39 | 3.74 | 1.99 | 1.42 | 7.96 | 0.32 |

| Measured Antigen | Exposure History | No. of Sera | Median Fold Breadth Gain | Std Dev | 2.5% Quantile | 97.5% Quantile | Standard Error |
|---|---|---|---|---|---|---|---|
| Omicron_BA.1 | 2-Dose Omicron BA.1/BA.1.1 PVI | 11 | 38.7593646 | 31.3 | 8.10899 | 106.3317 | 9.431909 |
| Omicron_BA.1 | 3-Dose Omicron BA.1/BA.1.1 PVI | 10 | 40.8737894 | 10.5 | 29.577 | 59.92547 | 3.310173 |
| Omicron_BA.1 | 3 Dose Vaccine Only | 39 | 19.0428004 | 11.2 | 8.30111 | 48.03459 | 1.791968 |
| Omicron_BA.2 | 2-Dose Omicron BA.1/BA.1.1 PVI | 11 | 17.0831321 | 8.97 | 4.98513 | 34.72302 | 2.703377 |
| Omicron_BA.2 | 3-Dose Omicron BA.1/BA.1.1 PVI | 10 | 17.2744759 | 3.54 | 13.5584 | 24.76249 | 1.1204 |
| Omicron_BA.2 | 3 Dose Vaccine Only | 39 | 12.3011188 | 6.95 | 3.91527 | 28.31128 | 1.112644 |
| Omicron_BA.2.12.1 | 2-Dose Omicron BA.1/BA.1.1 PVI | 11 | 12.5332508 | 7.3 | 2.61599 | 24.95442 | 2.200086 |
| Omicron_BA.2.12.1 | 3-Dose Omicron BA.1/BA.1.1 PVI | 10 | 11.047997 | 2.76 | 7.67431 | 16.56612 | 0.87307 |
| Omicron_BA.2.12.1 | 3 Dose Vaccine Only | 39 | 7.18766883 | 4.09 | 0.58812 | 15.20423 | 0.654776 |
| Omicron_BA.4_BA.5 | 2-Dose Omicron BA.1/BA.1.1 PVI | 11 | 5.82447672 | 4.43 | 0.75544 | 14.70707 | 1.336792 |
| Omicron_BA.4_BA.5 | 3-Dose Omicron BA.1/BA.1.1 PVI | 10 | 5.89732187 | 1.99 | 1.88801 | 7.782728 | 0.630663 |
| Omicron_BA.4_BA.5 | 3 Dose Vaccine Only | 39 | 3.10406937 | 2.81 | 1.35345 | 7.692711 | 0.450164 |
| Omicron_BA.1.1 | 2-Dose Omicron BA.1/BA.1.1 PVI | 11 | 62.9774641 | 33.1 | 7.65224 | 107.3031 | 9.989195 |
| Omicron_BA.1.1 | 3-Dose Omicron BA.1/BA.1.1 PVI | 10 | 48.4354099 | 10.6 | 38.8104 | 72.18467 | 3.358569 |

**Table S8: Significance values for breadth gain comparisons**

(Bolded comparisons have p-values lower than 0.05)

| Measured Antigen | Broader Exposure | Narrower Exposure | Test Stat | P Value | Stat Low CI | Stat Up CI |
|---|---|---|---|---|---|---|
| WT_D614G | 3-Dose Omicron BA.1/BA.1.1 PVI | 3 Dose Vaccine Only | 200 | 0.3242911 | 0 | Inf |
| WT_D614G | 3-Dose Omicron BA.1/BA.1.1 PVI | 2-Dose Omicron BA.1/BA.1.1 PVI | 65 | 0.09465659 | 0 | Inf |
| WT_D614G | 2-Dose Omicron BA.1/BA.1.1 PVI | 3 Dose Vaccine Only | 181 | 0.9734615 | -2.87E-05 | Inf |
| **Delta_B.1.617.2** | **3-Dose Omicron BA.1/BA.1.1 PVI** | **3 Dose Vaccine Only** | **386** | **1.15E-06** | **0.655286** | **Inf** |
| Delta_B.1.617.2 | 3-Dose Omicron BA.1/BA.1.1 PVI | 2-Dose Omicron BA.1/BA.1.1 PVI | 71 | 0.13745355 | -0.14221 | Inf |
| **Delta_B.1.617.2** | **2-Dose Omicron BA.1/BA.1.1 PVI** | **3 Dose Vaccine Only** | **414** | **1.58E-06** | **0.394052** | **Inf** |
| Mu_B.1.621 | 3-Dose Omicron BA.1/BA.1.1 PVI | 3 Dose Vaccine Only | 229 | 0.20297757 | -0.39143 | Inf |
| Mu_B.1.621 | 3-Dose Omicron BA.1/BA.1.1 PVI | 2-Dose Omicron BA.1/BA.1.1 PVI | 56 | 0.48595657 | -0.75987 | Inf |
| Mu_B.1.621 | 2-Dose Omicron BA.1/BA.1.1 PVI | 3 Dose Vaccine Only | 246 | 0.23391838 | -0.46918 | Inf |
| Beta_B.1.351 | 3-Dose Omicron BA.1/BA.1.1 PVI | 3 Dose Vaccine Only | 185 | 0.60275051 | -1.51375 | Inf |
| Beta_B.1.351 | 3-Dose Omicron BA.1/BA.1.1 PVI | 2-Dose Omicron BA.1/BA.1.1 PVI | 48 | 0.70129687 | -1.18221 | Inf |
| Beta_B.1.351 | 2-Dose Omicron BA.1/BA.1.1 PVI | 3 Dose Vaccine Only | 208 | 0.56510933 | -1.39419 | Inf |

| Measured Antigen | Broader Exposure | Narrower Exposure | Test Stat | P Value | Stat Low CI | Stat Up CI |
|---|---|---|---|---|---|---|
| **Omicron_BA.4_BA.5** | **3-Dose Omicron BA.1/BA.1.1 PVI** | **3 Dose Vaccine Only** | **272** | **0.02886573** | **0.57307** | **Inf** |
| Omicron_BA.4_BA.5 | 3-Dose Omicron BA.1/BA.1.1 PVI | 2-Dose Omicron BA.1/BA.1.1 PVI | 49 | 0.67642183 | -3.26783 | Inf |
| **Omicron_BA.4_BA.5** | **2-Dose Omicron BA.1/BA.1.1 PVI** | **3 Dose Vaccine Only** | 295 | **0.03049576** | **0.460743** | **Inf** |
| **Omicron_BA.2** | **3-Dose Omicron BA.1/BA.1.1 PVI** | **3 Dose Vaccine Only** | 293 | **0.00778853** | **2.067469** | **Inf** |
| Omicron_BA.2 | 3-Dose Omicron BA.1/BA.1.1 PVI | 2-Dose Omicron BA.1/BA.1.1 PVI | 56 | 0.48595657 | -4.08854 | Inf |
| **Omicron_BA.2** | **2-Dose Omicron BA.1/BA.1.1 PVI** | **3 Dose Vaccine Only** | **301** | **0.0220009** | **1.411535** | **Inf** |
| **Omicron_BA.2.12.1** | **3-Dose Omicron BA.1/BA.1.1 PVI** | **3 Dose Vaccine Only** | **316** | **0.00139838** | **1.737903** | **Inf** |
| Omicron_BA.2.12.1 | 3-Dose Omicron BA.1/BA.1.1 PVI | 2-Dose Omicron BA.1/BA.1.1 PVI | 39 | 0.87736094 | -6.52204 | Inf |
| **Omicron_BA.2.12.1** | **2-Dose Omicron BA.1/BA.1.1 PVI** | **3 Dose Vaccine Only** | **337** | **0.00213716** | **2.708569** | **Inf** |
| **Omicron_BA.1** | **3-Dose Omicron BA.1/BA.1.1 PVI** | **3 Dose Vaccine Only** | **363** | **1.63E-05** | **16.37823** | **Inf** |
| Omicron_BA.1 | 3-Dose Omicron BA.1/BA.1.1 PVI | 2-Dose Omicron BA.1/BA.1.1 PVI | 55 | 0.51404343 | -16.8562 | Inf |
| **Omicron_BA.1** | **2-Dose Omicron BA.1/BA.1.1 PVI** | **3 Dose Vaccine Only** | **345** | **0.00116522** | **11.68125** | **Inf** |
| Omicron_BA.1.1 | 3-Dose Omicron BA.1/BA.1.1 PVI | 2-Dose Omicron BA.1/BA.1.1 PVI | 48 | 0.70129687 | -27.3662 | Inf |

**Table S9: Slope and Peak Location for Individual Antibody Landscapes**

| Exposure History | RECORD_ID | Cone Peak X Coord | Cone Peak Y Coord | Cone Slope |
|---|---|---|---|---|
| 3 Dose Vaccine Only | 1009 | -2.234571435 | 1.305183215 | 0.393543202 |
| 3 Dose Vaccine Only | 1012 | -1.538545488 | 1.885725339 | 0.459436789 |
| 3 Dose Vaccine Only | 1013 | -3.764331536 | 1.398561998 | 0.255213906 |
| 3 Dose Vaccine Only | 1018 | -1.903423786 | 2.444181951 | 0.450580611 |
| 3 Dose Vaccine Only | 1021 | -2.016754846 | 1.532996781 | 0.578235008 |
| 3 Dose Vaccine Only | 1026 | -2.274626285 | 2.12991451 | 0.434504268 |
| 3 Dose Vaccine Only | 1027 | -1.358131089 | 1.758091423 | 0.512011971 |
| 3 Dose Vaccine Only | 1029 | -2.227493152 | 1.577981324 | 0.491516828 |
| 3 Dose Vaccine Only | 1031 | -2.874562598 | 2.059062376 | 0.255906208 |
| 3 Dose Vaccine Only | 1032 | -2.144557765 | 1.625794771 | 0.39182111 |
| 3 Dose Vaccine Only | 1037 | -0.796959015 | 2.890020707 | 0.383170571 |
| 3 Dose Vaccine Only | 1039 | -1.240451449 | 2.179803449 | 0.553607988 |
| 3 Dose Vaccine Only | 1043 | -1.408899264 | 1.699739244 | 0.546262723 |
| 3 Dose Vaccine Only | 1060 | -2.09410053 | 2.082915253 | 0.359706363 |
| 3 Dose Vaccine Only | 1074 | -2.833411974 | 1.320033197 | 0.316253181 |
| 3 Dose Vaccine Only | 1076 | -1.199941913 | 2.49086648 | 0.454757494 |
| 3 Dose Vaccine Only | 1078 | -2.148029665 | 1.690572344 | 0.437972805 |
| 3 Dose Vaccine Only | 1079 | -3.283442632 | 0.826959509 | 0.326190606 |
| 3 Dose Vaccine Only | 1082 | -1.759280788 | 1.655158957 | 0.527723863 |
| 3 Dose Vaccine Only | 1103 | -2.853188611 | 1.622334486 | 0.425622426 |
| 3 Dose Vaccine Only | 1113 | -2.142498445 | 1.323119727 | 0.393842798 |
| 3 Dose Vaccine Only | 1126 | -1.840047168 | 1.658686652 | 0.417468602 |
| 3 Dose Vaccine Only | 1137 | -0.896503487 | 2.497227404 | 0.474622941 |
| 3 Dose Vaccine Only | 1150 | -1.553991662 | 2.361255188 | 0.452735856 |
| 3 Dose Vaccine Only | 1153 | -1.971358365 | 2.658891414 | 0.399509443 |
| 3 Dose Vaccine Only | 1156 | -1.824630217 | 2.337446299 | 0.405180622 |
| 3 Dose Vaccine Only | 1160 | -2.484890802 | 0.895753825 | 0.488872145 |
| 3 Dose Vaccine Only | 1166 | -1.663649266 | 0.639071374 | 0.505807478 |
| 3 Dose Vaccine Only | 1172 | -1.521308691 | 1.102692766 | 0.619519741 |
| 3 Dose Vaccine Only | 1173 | -1.963067301 | 1.154955937 | 0.541419819 |
| 3 Dose Vaccine Only | 1180 | -1.867613678 | 2.099370046 | 0.429335136 |
| 3 Dose Vaccine Only | 1209 | -1.02767303 | 2.61278104 | 0.381233654 |
| 3 Dose Vaccine Only | 1215 | -2.441673845 | 2.405299416 | 0.294995553 |
| 3 Dose Vaccine Only | 1216 | -0.183128945 | 3.336613664 | 0.26270095 |
| 3 Dose Vaccine Only | 1221 | -1.025796527 | 3.080732552 | 0.385018872 |
| 3 Dose Vaccine Only | 1229 | -2.070676499 | 0.710100186 | 0.861094318 |
| 3 Dose Vaccine Only | 1244 | -1.991742263 | 1.207060325 | 0.712158321 |
| 3 Dose Vaccine Only | 1255 | -1.462972005 | 1.961360838 | 0.511233743 |
| 3 Dose Vaccine Only | 1274 | -1.237332418 | 2.54898911 | 0.382689339 |

**Table S10: Summary statistics of cone slope for individual landscapes grouped by exposure history**

| Exposure History | # of Sera | Median Cone Slope | Std Dev | 2.5% Quantile | 97.5% Quantile | Std Error |
|---|---|---|---|---|---|---|
| 2-Dose Omicron BA.1/BA.1.1 PVI | 11 | 0.280229166 | 0.182 | 0.12839666 | 0.698305257 | 0.0549 |
| 3-Dose Omicron BA.1/BA.1.1 PVI | 10 | 0.301508148 | 0.0383 | 0.22396893 | 0.330718312 | 0.0121 |
| 3 Dose Vaccine Only | 39 | 0.434504268 | 0.1195 | 0.25587159 | 0.719605121 | 0.0191 |

**Fig. S1**

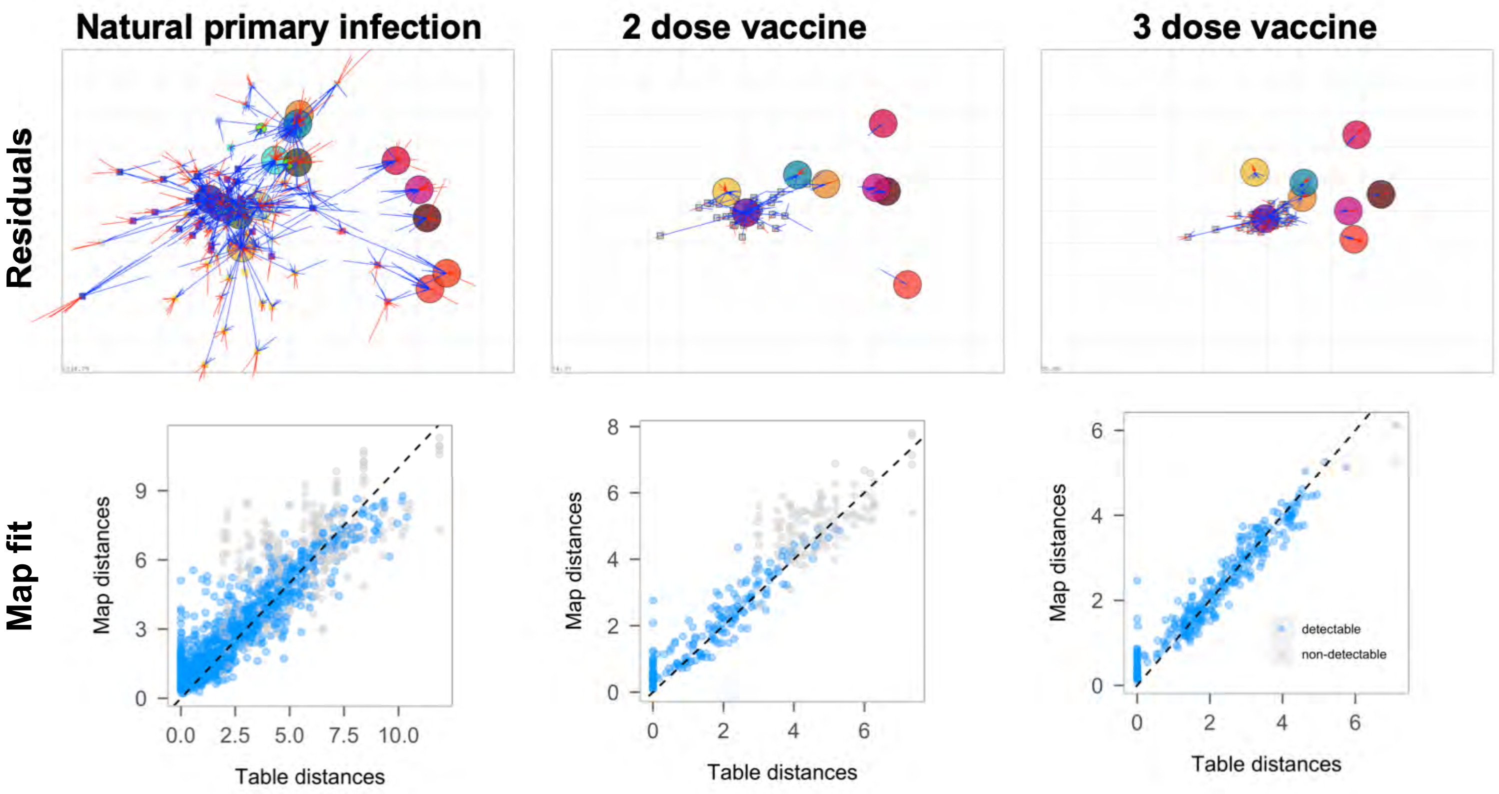


Dimensionality

Natural primary infection

| Dim. | Detectable Mean RMSE | Detectable Var. RMSE | Undetectable Mean RMSE | Undetectable Var. RMSE |
|---|---|---|---|---|
| 1 | 1.54 | 0.05 | 1.95 | 0.09 |
| 2 | 1.36 | 0.04 | 1.73 | 0.07 |
| 3 | 1.34 | 0.04 | 1.61 | 0.05 |
| 4 | 1.36 | 0.04 | 1.58 | 0.05 |
| 5 | 1.36 | 0.04 | 1.57 | 0.05 |

2 dose vaccine

| Dim. | Detectable Mean RMSE | Detectable Var. RMSE | Undetectable Mean RMSE | Undetectable Var. RMSE |
|---|---|---|---|---|
| 1 | 1.28 | 0.28 | 1.01 | 0.06 |
| 2 | 1.39 | 0.44 | 0.98 | 0.18 |
| 3 | 1.40 | 0.44 | 0.92 | 0.07 |
| 4 | 1.39 | 0.38 | 0.92 | 0.10 |
| 5 | 1.40 | 0.43 | 0.92 | 0.08 |

3 dose vaccine

| Dim. | Detectable Mean RMSE | Detectable Var. RMSE | Undetectable Mean RMSE | Undetectable Var. RMSE |
|---|---|---|---|---|
| 1 | 1.00 | 0.04 | 2.08 | 0.62 |
| 2 | 1.06 | 0.05 | 1.91 | 0.21 |
| 3 | 1.07 | 0.05 | 1.89 | 0.21 |
| 4 | 1.07 | 0.05 | 1.90 | 0.21 |
| 5 | 1.07 | 0.05 | 1.90 | 0.21 |

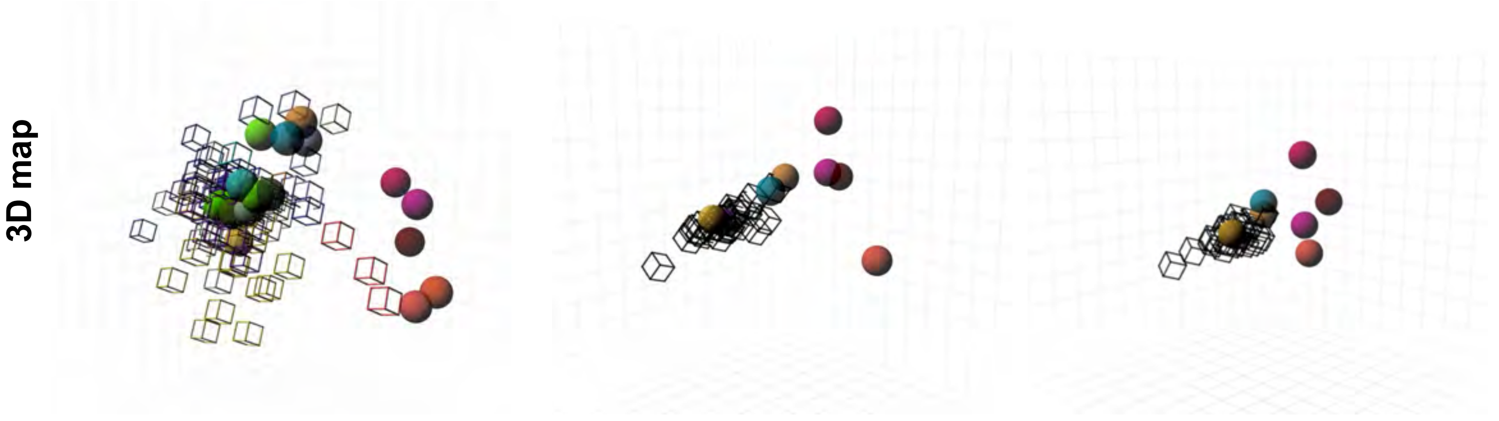

**Fig. S2**

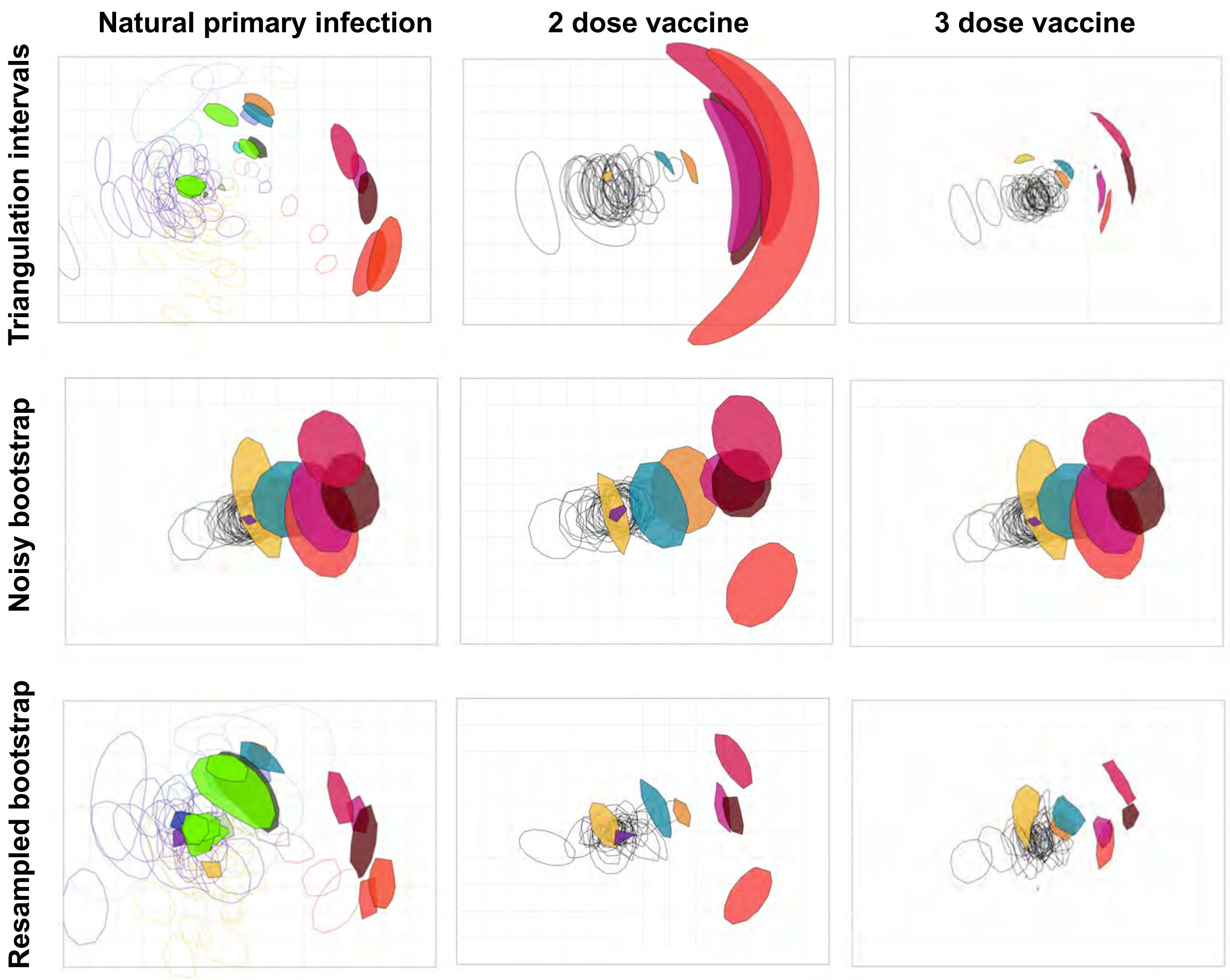

Natural primary infection
2 dose vaccine
3 dose vaccine
Triangulation intervals
Noisy bootstrap
Resampled bootstrap

Fig. S3

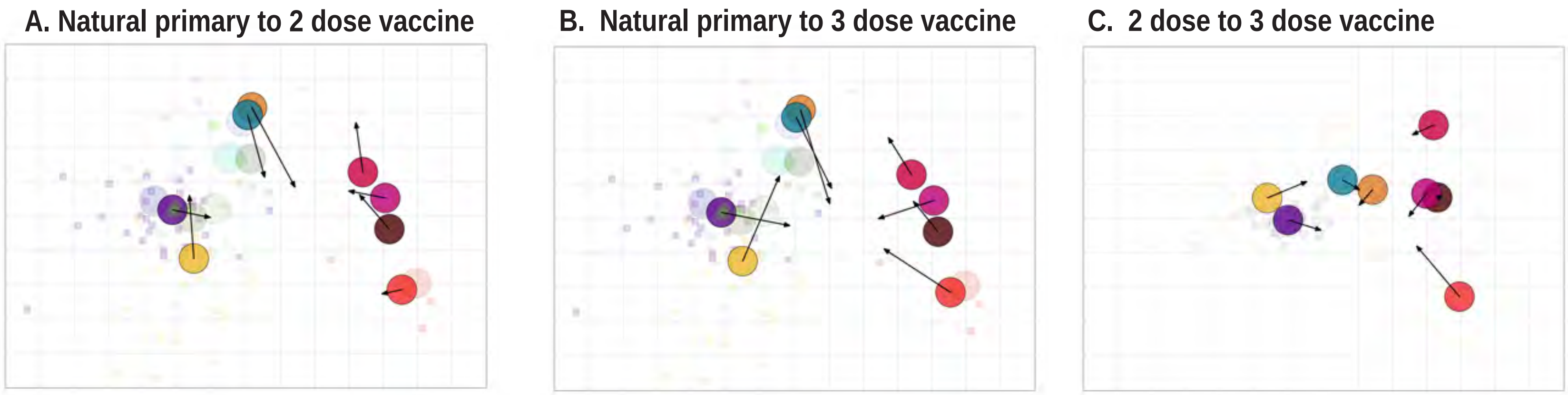

A. Natural primary to 2 dose vaccine
B. Natural primary to 3 dose vaccine
C. 2 dose to 3 dose vaccine

### Supplementary Figure legends

**Fig. S1. Evaluation of goodness of fit and dimensionality for antigenic maps made with primary infection antisera (column 1), two dose vaccine sera (column 2), and three dose vaccine sera (column 3).** Sera are shown as small colored squares, viruses as large circles. The grid corresponds to a two-fold dilution in the neutralization assay. Row 1 shows the antigenic map with error lines. The distance between the ends of error lines indicates the measured titer: red lines indicate that the map distance is less than measured based on the titers, blue lines when the map distance is greater than measured. Row 2 shows the difference between the table distance (estimated from the measured titer) and the fitted map distance. The dotted horizontal line shows what would be perfect a perfect fit of the data. Row 3 shows the results of dimensionality testing. Cross-validation (excluding 10% of titers as a test set in 100 independent repeats) was used to determine the optimal number of dimensions. Lower root mean squared error (RMSE) for both detectable titers (above the assay limit of detection) and undetectable (below the assay limit of detection) indicate the optimal number of dimensions for fitting the antigenic map. Row 4 shows antigenic maps made in three dimensions.

**Fig. S2. Evaluation of robustness in positioning for viruses and sera on antigenic maps made with primary infection antisera (column 1), two dose vaccine sera (column 2), and three dose vaccine sera (column 3).** Sera are shown as open shapes, viruses as colored shapes. The grid corresponds to a two-fold dilution in the neutralization assay. Row 1 shows triangulation/coordination confidence intervals, indicating confidence in positioning of points.

Each shape marks the area that the point can occupy before increasing the total map error by more than 1 antigenic unit. Row 2 shows bootstrapped maps considering titer error for the neutralization assay. The shapes correspond to the positions of points on resampled maps assuming titers have random noise added with the measured assay standard deviation of log2 0.29 (1.2-fold). Row 3 shows confidence in coordination of points following bootstrapping of the sera and viruses.

**Fig. S3.** Comparison of virus positions between antigenic maps. Arrows point to virus positions from one map to another. Sera are shown as small squares, viruses as colored circles. The grid corresponds to a two-fold dilution in the neutralization assay.
